# Nuclear Location Bias of HCAR1 Drives Cancer Malignancy through Numerous Routes

**DOI:** 10.1101/2022.07.25.501445

**Authors:** Mohammad Ali Mohammad Nezhady, Gael Cagnone, Emmanuel Bajon, Prabhas Chaudhari, Monir Modaresinejad, Pierre Hardy, Damien Maggiorani, Christiane Quiniou, Jean-Sébastien Joyal, Christian Beauséjour, Sylvain Chemtob

**Affiliations:** Program in Molecular Biology, Faculty of Medicine, Université de Montréal, Montreal, QC, Canada; Research Center of Centre Hospitalier Universitaire Sainte-Justine, Montreal, QC H3T 1C5, Canada; Department of Experimental Medicine, McGill University, Montréal, Canada; Program in Biomedical Sciences, Faculty of Medicine, Université de Montréal, Montreal, QC, Canada; Department of Pharmacology, Université de Montréal, Montreal, Quebec, Canada

**Keywords:** HCAR1, Lactate, GPCR, Location bias, Nucleus, Signaling, Cancer, Warburg effect

## Abstract

The involvement of G-Protein-Coupled Receptors’ (GPCR) location bias in diverse cellular functions and their misregulation in pathology is an underexplored territory. HCAR1, a GPCR for lactate is linked to cancer progression, mainly due to Warburg effect, but its mechanism of action remains elusive. Here, we show HCAR1 has a nuclear localization, capable of signaling intranuclearly to induce nuclear-ERK and AKT phosphorylation concomitant with higher cancer cell proliferation and survival. We determine its nuclear interactome, proving its involvement in protein-translation and DNA-damage repair. Nuclear HCAR1 (N-HCAR1) directly interacts with chromatin/DNA promoting expression of genes involved in cellular migration. Notably, we show N-HCAR1 particularly regulates a broader transcriptomic signature than its PM counterpart, emphasizing on the facts that functional output of N-HCAR1 is larger than PM localized HCAR1. Our study presents several unprecedented processes by which a GPCR through location-biased activity regulate various cellular functions and how cancer cells exploit these.

## Introduction

Hydroxycarboxylic acid receptor 1 (HCAR1), a G-Protein Coupled Receptor (GPCR) also known as GPR81, initially was shown to inhibit lipolysis (Ge *et al*., 2008). Discovery of lactate as its ligand (Liu *et al*., 2009) led to an intensive, yet ongoing research to attribute many effects of lactate to HCAR1. In this context, major focus of recent efforts in the field are placed on cancer studies due to increased level of lactate production in most tumors because of the Warburg effect (Heiden, Cantley and Thompson, 2009),(Zhao *et al*., 2020),(Brown *et al*., 2020),(Brown and Ganapathy, 2020). Despite high energy demand in cancer cells for rapid proliferation, these cells switch to glycolysis as the major source of energy production causing accumulation of the end product lactate (Liberti and Locasale, 2016); a consequence of the Warburg effect is a substantial increase in the concentration of lactate in the tumor niche (up to 40 mM)(Sun *et al*., 2017). Although several hypotheses have been proposed to explain this phenomenon, a solid explanation for this rather paradoxical metabolic switch remains to be described (Devic, 2016).

Lactate acting through HCAR1 exerts diverse effects in promoting cancer malignancy. *HCAR1* is overexpressed in numerous cancer cell lines and resected tumors from patients (Roland *et al*., 2014),(Lee *et al*., 2016), (Stäubert, Broom and Nordström, 2015). Orthotopic xenografts of pancreatic and breast cancer cells depleted in HCAR1 have markedly lower growth and proliferation rates, metastasis, angiogenesis and survival compared to endogenously *HCAR1* expressed cancer cell xenografts (Roland *et al*., 2014),(Lee *et al*., 2016). HCAR1 stimulation with lactate directly induces expression of angiogenic factors such as *Vascular endothelial growth factor (VEGF)* and *Amphiregulin (AREG)* in normal and malignant conditions, respectively (Morland *et al*., 2017),(Lee *et al*., 2016). *In vitro*, lactate through HCAR1 promotes DNA damage repair capacity, thus conferring resistance to DNA damage-inducing chemotherapeutic agents. Interestingly, although HCAR1 is considered a cell surface receptor, its action on promoting DNA damage repair was restricted when intracellular lactate uptake was inhibited, meaning that extracellular lactate concentration required to elicit cell surface HCAR1 signaling was not determinant for the receptor function (Wagner, Ciszewski and Kania, 2015),(Wagner, Kania and Ciszewski, 2017). In addition, HCAR1 mitigates cancer immune evasion, another hallmark of cancer malignancy. Lactate via HCAR1 induces expression of *PD-L1* in tumor cells with consequent reduced interferon (IFN)-γ production resulting in potential immune escape (Feng *et al*., 2017). HCAR1 immune evasive roles are not limited to tumor cells, HCAR1 in immune cells also contributes to this effect (Raychaudhuri *et al*., 2019). IFN-α production was abolished in dendritic cells upon HCAR1 activation with lactate, which in turn enhanced tumor evasion and growth; interestingly, inhibition of intra-cellular lactate uptake again diminished effects of HCAR1 (Raychaudhuri *et al*., 2019). However, mechanisms to explain multidimensional involvement of HCAR1 in cancer biology is lacking; and an intracellular mode of action should be accounted for.

HCAR1 is considered a plasma membrane receptor which signals intracellularly. Yet, there is an emergence in discovery of functional intracellular GPCRs. Every membranous organelle has been shown to harbor active GPCRs, either as a primary site of localization or as a result of plasma membrane translocation upon ligand binding (Jong, Harmon and O’Malley, 2018),(Mohammad Nezhady, Rivera and Chemtob, 2020). The differential activity of a GPCR from these intracellular organelles as opposed to their signaling output from plasma membrane is generally referred to as location bias (Mohammad Nezhady, Rivera and Chemtob, 2020). Even the spatial cellular coordination from where a GPCR activates the same signaling pathway could lead to differential transcriptional output demonstrating the significance of location bias (Tsvetanova and von Zastrow, 2014). Moreover some GPCRs are primarily resident within the nucleus (Bhosle *et al*., 2016) or endoplasmic reticulum (Revankar *et al*., 2005), (Sanchez *et al*., 2022). These facts emphasize considering subcellular localization of GPCRs when investigating their biology.

Here, we show HCAR1 has a nuclear localization, we decipher its topology on the nuclear membranes, and reveal its interactome and function. We show that nuclear HCAR1 (N-HCAR1) is capable of initiating intranuclear signaling. Our bottom-up high-throughput omics studies demonstrate that N-HCAR1 promotes various processes by different mechanisms including protein translation and DNA damage repair involved in cell proliferation and propagation. Notably, we show N-HCAR1 regulates a broader transcriptomic signature than its plasma membrane counterpart, emphasizing on the facts that N-HCAR1 functional output is larger than plasma membrane localized HCAR1, and highlights the importance of receptor location bias. Furthermore, we validate these effects *in vivo* providing a mechanistic view to various roles of HCAR1 in promoting cancer malignancy.

## Results

### HCAR1 has a nuclear localization pattern mediated through the 3^rd^ intracellular loop domain and S305 phosphorylation

We generated stable HeLa cell lines expressing either C-terminal or N-terminal Flag-tagged HCAR1, allowing for enhanced localization as described in Methods. HCAR1 nuclear localization was ascertained using various methods. Complete separation of nucleus upon biochemical cell fractionation revealed abundant HCAR1 at the nucleus, as well as in the cytoplasm (Fig. 1a). Immunofluorescent staining with confocal microscopy using Lamin B1 as inner nuclear membrane marker exhibited clear HCAR1 colocalization with Lamin B1 in intact cells and isolated nuclei; strikingly, it was also detectable inside the nucleus (Fig. 1b,c; Supp Fig. 1a,d; Fig. 2a). Immunogold staining on electron microscopy confirmed nuclear envelope and intranuclear HCAR1 distribution (Fig. 1d; Supp Fig. 1e-g); nearly one-third of cellular HCAR1 was localized at the nucleus in unstimulated cells (Fig. 1e). Nuclear localization of HCAR1 was also detected in U251MG and A549 HCAR1-transfected cells (Supp Fig. 2a,d).

**Fig. 1:**
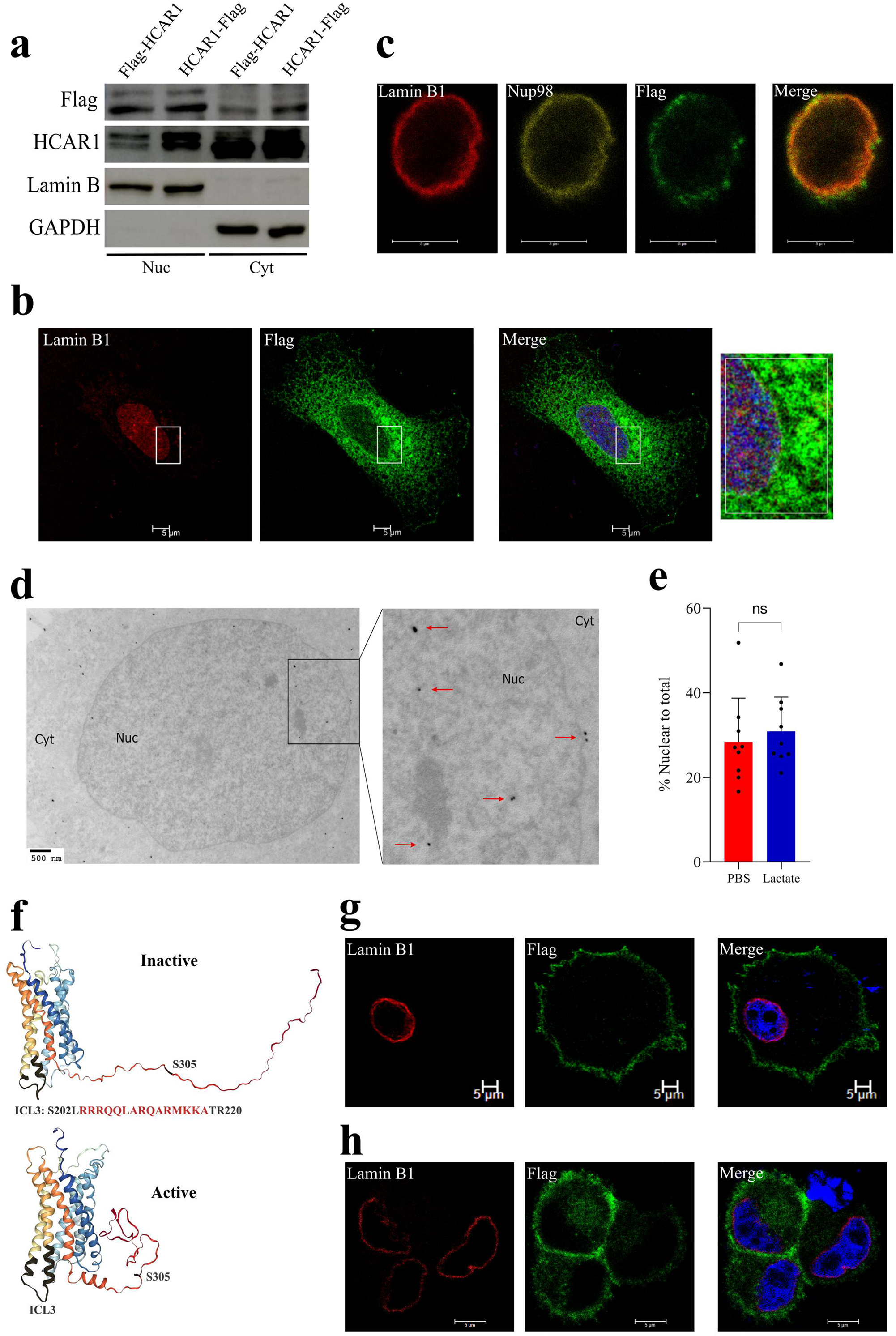
HCAR1 is present in the nucleus and ICL3 and S305 phosphorylation are responsible for this localization pattern. **a)** Western blot analysis of fractionated cells transfected with C & N-terminally Flag-tagged HCAR1. Lamin B and GAPDH were used to confirm pure isolation of nuclei. **b-c)** Confocal imaging of C-terminally flag-tagged HCAR1 whole cells **(b)** or isolated nuclei **(c). d)** TEM graphs from C-terminally flag-tagged HCAR1. **e)** Quantification HCAR1 from TEM images of PBS and Lactate treated cells (10mM for 1h). **f)** 3D modeling of HCAR1 in inactive and active conformations by GPCRM. The black highlights indicate the spanning regions for ICL3 domain and S305. **g-h)** Confocal imaging of C-terminally flag-tagged HCAR1 with ICL3 deletion **(g)** and S305A mutation **(h)**. Notice the cytoplasmic signal of HCAR1 in S305A.

**Fig. 2:**
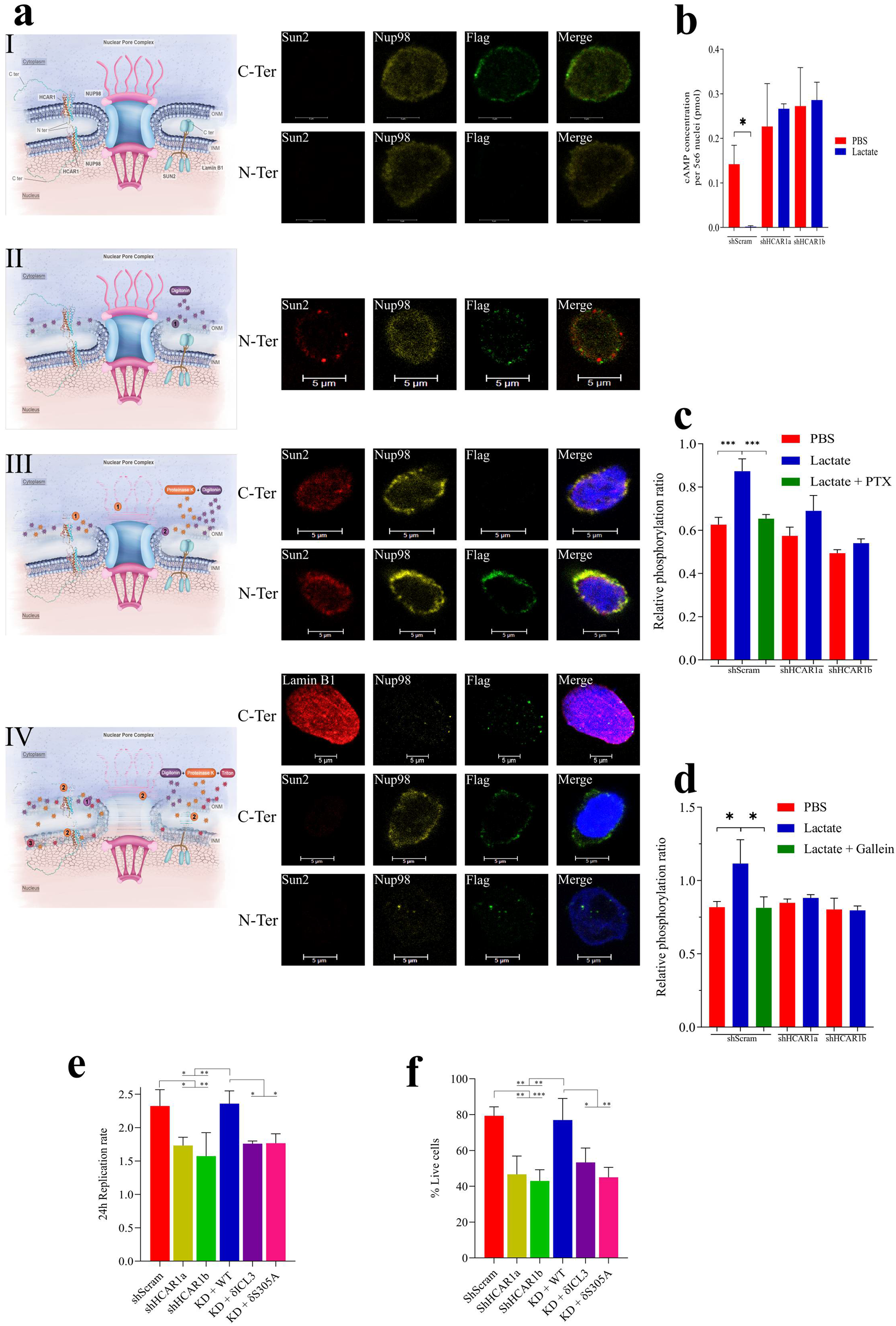
Intranuclear signaling of HCAR1 activates nuclear ERK and AKT effectors leading to cellular proliferation and survival. **a)** Confocal images of nuclei isolated from cells expressing C-ter on N-ter flag-tagged HCAR1. **(I)** intact nuclei, **(II)** ONM permeabilized nuclei with intact INM, **(III)** surface protein digested nuclei with ONM permeabilization and intact INM, **(IV)** ONM permeabilized nuclei with intact INM was treated with PK to digest proteins on the ONM and lumen, and after washing PK, nuclei were treated with triton to permeabilize INM. Notice loss of Sun2 indicating digestion of luminal proteins. **b)** cAMP level in isolated nuclei from scrambled shRNA or two different HCAR1 KD cells with PBS or lactate treatment (10mM for 15min). The cAMP concentration is presented in picomole per 5 million nuclei. **c-d)** ELISA analysis of ERK **(c)** and AKT **(d)** phosphorylation rates in isolated nuclei from scrambled shRNA or two different HCAR1 KD cells with PBS or lactate treatment (10mM for 15min). PTX or Gallein treatment of scrambled cells were performed prior to nuclei isolation. **e)** Cell proliferation rate in scrambled shRNA, two different HCAR1 KD cells, WT-rescue and nuclear KD cell lines. **f)** Cellular survival rate in 5FU treated cells. Data are mean ± s.d. from n≥3 biological replicates. Analysis of Variance (ANOVA) was followed by Bonferroni post hoc correction test with * *P* < 0.05, ** *P* < 0.01, \*\*\**P* < 0.0001 significance levels.

Treatment of cells with lactate did not alter the nuclear ratio of HCAR1, suggesting there is no translocation from plasma membrane to the nucleus upon ligand stimulation (Fig. 1e; Supp Fig. 1h). To further investigate this possibility, we devised a pulse chase study using Fluorogen Activating Peptide (FAP) technology utilizing cell impermeable fluorogen (Fisher *et al*., 2014). Activation of the receptor with lactate, triggered HCAR1 internalization within 5 min; HCAR1-containing endosomes were tracked in the cytoplasm for up to 40 min, after which it was no longer detected (either because of recycling to the plasma membrane or endosomal degradation; Supp Fig. 3a-d). Although nuclear localization of HCAR1 from plasma membrane was not observed, the chimeric receptor was hitherto present in the nucleus prior to lactate stimulation (Supp Fig. 3e). This observation along with the electron microscopy data indicates that there is a *de facto* nuclear pool of HCAR1 present in the cells.

In an attempt to determine HCAR1 domains necessary for nuclear localization we analyzed the 3D model of the receptor (Fig. 1f) and found that although there is no classical nuclear localization signal (NLS) in HCAR1 sequence, there are predicated bipartite NLS in intracellular loop 3 (ICL3), and C-terminus of the receptor (Kosugi *et al*., 2009). Truncation in ICL3 totally abolished nuclear localization as well as cytoplasmic staining (Fig. 1g; Supp Fig. 2b,e); efforts to pinpoint precise amino acid sequences necessary for nuclear localization were not fruitful. A phosphorylation site in the NLS of the C-terminus (Hornbeck *et al*., 2015), prompted us to determine its potential role in localization. Single point substitution of S305 to alanine at the C-terminus led to exclusion of HCAR1 from the nucleus (but retained cytoplasmic staining; Fig. 1h; Supp Fig. 2c,f). These findings suggest a scaffolding role of ICL3 and post translational phosphorylation of S305 required for HCAR1 nuclear localization.

### Nuclear HCAR1 is functional at inner nuclear membrane, signaling for proliferation and survival

To better understand the function of HCAR1 through downstream signaling cascade, we first determined the topology of HCAR1 on both outer and inner nuclear membranes (ONM, INM, respectively). Immunofluorescence staining of intact (non-permeabilized) nuclei isolated from HCAR1 C-terminal Flag-tagged cells indicates that the C-terminus of the receptor is oriented towards the cytoplasm on the ONM (Fig. 2aI). Whereas, intact nuclei from HCAR1 N-terminal Flag-tagged cells did not display any staining, consistent with suggestion that the N-terminus of the receptor points within the nuclear envelope (Fig. 2aI). To ascertain this orientation, we devised a protocol to selectively permeabilize the ONM, while keeping the INM intact. To distinguish the differential permeabilization of the ONM and INM, we used a combination of 3 proteins as markers located in different parts of the nuclear envelope: a) NUP98 as a component of the nuclear pore, traverses the pore from whole outer to inner layer of the nuclear envelope, making it detectable in all the conditions (Xu and Powers, 2009). b) SUN2 spans the INM and its C-terminus is located in the lumen of the nuclear envelope (Chang, Worman and Gundersen, 2015); the antibody used targets the C-terminal domain of SUN2 (as luminal marker). c) Lamin B1 located on the nuclear side of the INM. Using 0.0008% digitonin as a mild detergent (Ledet *et al*., 2021), we were able to selectively permeabilize the ONM leaving the INM intact (Supp Fig. 4a). Permeabilization of the ONM (allowing antibody access) enabled to detect immunofluorescent signal for N-terminus Flag-tagged HCAR1, consistent with its luminal nuclear envelope localization (Fig. 2aII). Furthermore, treatment with proteinase K (PK) to remove the cytoplasm-facing C-terminus of HCAR1, followed by permeabilization of the ONM, revealed absent immuno-positive signal for the C-terminus but preserved the signal for the nuclear envelope lumen-localized N-terminus (Fig. 2aIII). We then proceeded to permeabilize the ONM, followed by sequential treatment of nuclei with PK and permeabilization of the INM; under these conditions, we again observed HCAR1 C-terminus staining co-localized with Lamin B1 (Fig. 2aIV). While the N-terminus in this condition was only detected inside the nucleus (not at its envelope). Altogether, these experiments reveal that the C-terminus of HCAR1 at the ONM orients within the cytoplasm, while at the INM it has analogous conformation to that at the plasma membrane to putatively initiate signaling cascade into the nucleus, as nuclear envelope membranes are known to contain signaling machinery (Gobeil *et al*., 2006).

To further elucidate this intranuclear signaling, we isolated intact nuclei and stimulated them with lactate and measured nuclear cAMP level. Lactate treatment of nuclei isolated from wildtype (WT) HeLa cells significantly decreased cAMP level compared to vehicle treated cells (Fig. 2b). Whereas, nuclei isolated from two different HCAR1 knocked down (KD) cells using distinct shRNAs (Supp Fig. 4f,g), did not respond to lactate (Fig. 2b). Similarly, lactate induced nuclear ERK1/2 and AKT phosphorylation in WT cells that were also unresponsive after HCAR1 KD (Fig. 2c,d; Supp Fig. 4b); notably, ERK1/2 phosphorylation was G_iα_-dependent (inhibited by pertussis toxin), and AKT phosphorylation was G_βγ_-dependent (inhibited by gallein) (Fig. 2c,d; Supp Fig. 4c,d).

Since ERK1/2 and AKT modulate proliferation and survival in cancer cells (Maik-Rachline, Hacohen-Lev-Ran and Seger, 2019), (Martelli *et al*., 2012), we measured homeostatic cell proliferation rate and cell survival upon 5-Fluorouracil (5-FU) challenge in cells expressing or devoid of nuclear HCAR1; former include WT HCAR1-expressing and HCAR1 KD cells rescued with RNAi resistant WT *HCAR1* (referred to as WT rescue cells; Supp Fig. 4e-g); the latter include HCAR1 KD cells rescued with RNAi resistant *HCAR1* containing δICL3 or δS305A mutations which excludes the receptor from the nucleus (referred to as N-HCAR1 KD cells; Supp Fig. 4e-g). HeLa cells harboring WT HCAR1 exhibited higher proliferation and survival rate compared to (un-rescued) HCAR1 KD and N-HCAR1 KD cells (Fig. 2e,f); nucleus excluded HCAR1 mutations caused similar effects in U251MG and A549 cancer cells (Supp Fig. 4h; Supp Fig. 5a,b). These data underscore the role of N-HCAR1 in cancer cell proliferation and survival.

### N-HCAR1 interacts with effectors involved in protein translation and DNA damage repair

Detection of HCAR1 inside the nucleus beside the NM prompted us to investigate spatiotemporal interactome of N-HCAR1. We used the Bio-ID system (Kim *et al*., 2016) to construct HCAR1-Bio-ID fusion protein which again revealed the expected intracellular localization of HA-tagged HCAR1 (Supp Fig. 6a,b). We isolated the nuclei of cells treated with or without lactate and purified their biotinylated proteome which was subjected for mass spectrometry (Fig. 3a). There was a clear distinction in the interactome of HCAR1 upon activation of the receptor (Fig. 3b; Supp Fig. 6c,d), suggesting that different conformations of N-HCAR1 participate in separate protein complexes (Fig. 3b). Surprisingly, the interactome of N-HCAR1 isolated from cells untreated with lactate was particularly enriched for proteins mediating tRNA aminoacylation involved in protein translation. While the interactome after lactate treatment was enriched for ribosomal regulatory processes (Fig. 3c; Supp Fig. 6e,f); this was validated on sucrose gradient ribosomal profiling which revealed lower content of non-polysomal ribosomes in HCAR1 KD and N-HCAR1 KD cells (Fig. 3d). Concordantly, protein translation was lower in HCAR1-KD and N-HCAR1 KD cells compared to cells with intact HCAR1 (Fig. 3e); similar observations on protein translation were made in U251MG and A549 cells (Supp Fig. 5c,d).

**Fig. 3:**
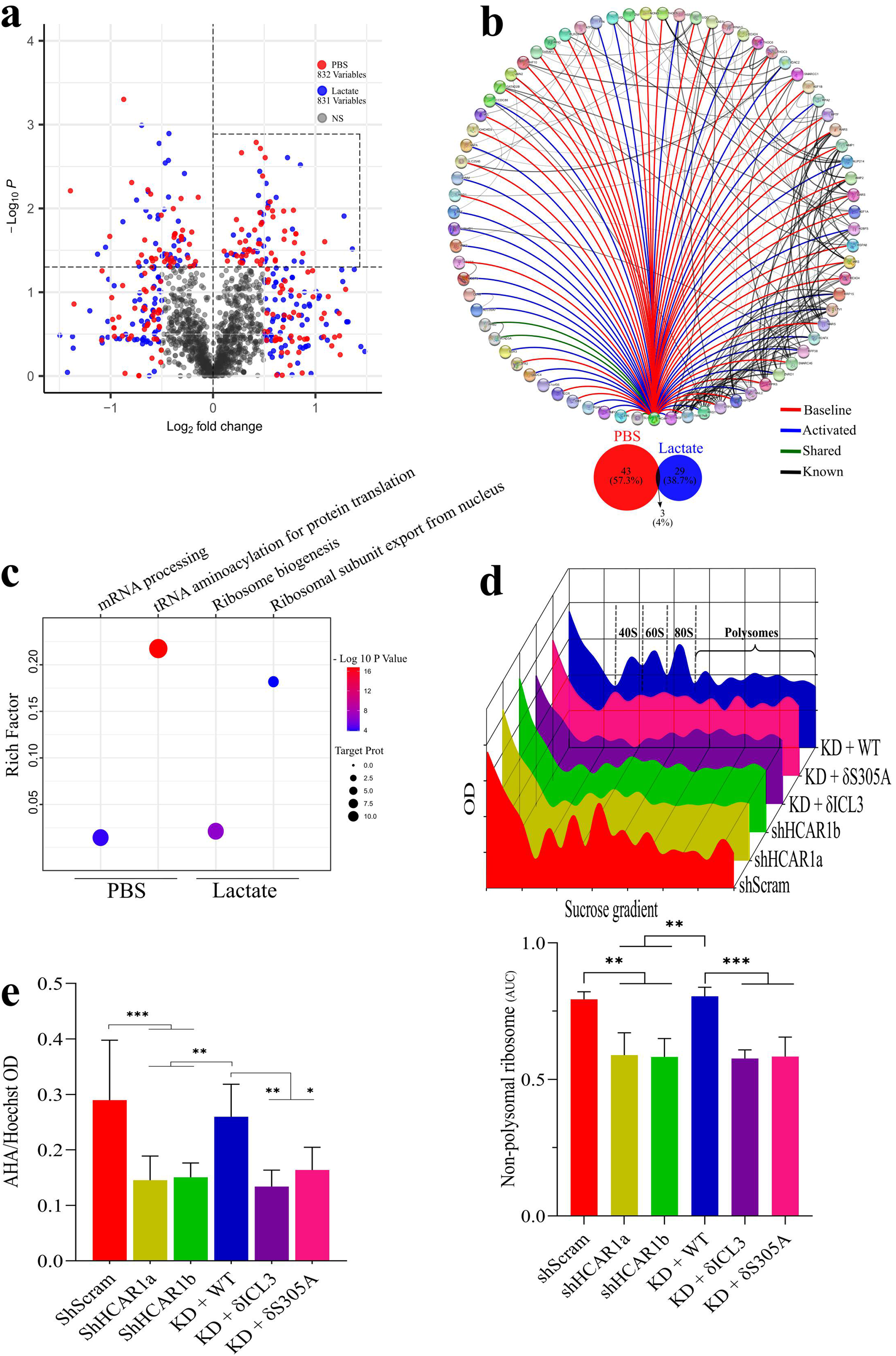
N-HCAR1 interactome is enriched for protein translational processes and it promotes protein translation rate. **a)** Volcano plot representing proteins enriched with HCAR1-BirA from Bio-ID experiment in isolated nuclei. Cells were treated with biotin and PBS or lactate (10mM for 24h) prior to nuclei isolation. **b)** Interactome map of N-HCAR1 in both PBS or lactate treated cells. The enriched proteins are selected based on both Log_2_ fold change and *p* value. Red lines indicate interaction of enriched proteins with HCAR1 when treated with PBS, blue lines indicate interactions with HCAR1 when treated with lactate, green lines indicate interaction in both cases, and black lines represents already established interactions based on Cytoscape. **c)** Enrichment dot plot for biological processes of proteins in panel b. **d)** Upper panel: Representative sucrose gradient ribosomal profiling for Scrambled shRNA, total and nuclear HCAR1 KD, and WT rescue cells. Lower panel: Normalized measurement of the upper panel for Area Under the Curve (AUC) of the monosomes (40S, 60S and 80S subunits). **f)** Protein translation rate with methionine incorporation rate measurement. Methionine incorporation rate (L-azidohomoalanine; AHA) was adjusted to the number of cells (Hoechst). Data are mean ± s.d. from n≥3 biological replicates. ANOVA was followed by Bonferroni post hoc correction test with * *P* < 0.05, ** *P* < 0.01, \*\*\**P* < 0.0001 significance levels.

In line with role of HCAR1 in cell proliferation and survival we detected components of DNA damage repair machinery in the interactome of N-HCAR1 in lactate treated and untreated cells, including the dominant DNA damage marker H2AX (Fig. 4a,b). Accordingly, irradiated WT and WT-HCAR1-rescued cells displayed lower DNA damage (γH2AX foci) upon recovery, compared to HCAR1-KD and N-HCAR1 KD cells, suggestive of limited DNA damage repair in nuclei devoid of HCAR1 (Fig. 4b). Together data show that N-HCAR1 interacts with classical signaling and non-signaling nuclear effectors that promote protein translation and DNA damage repair.

**Fig. 4:**
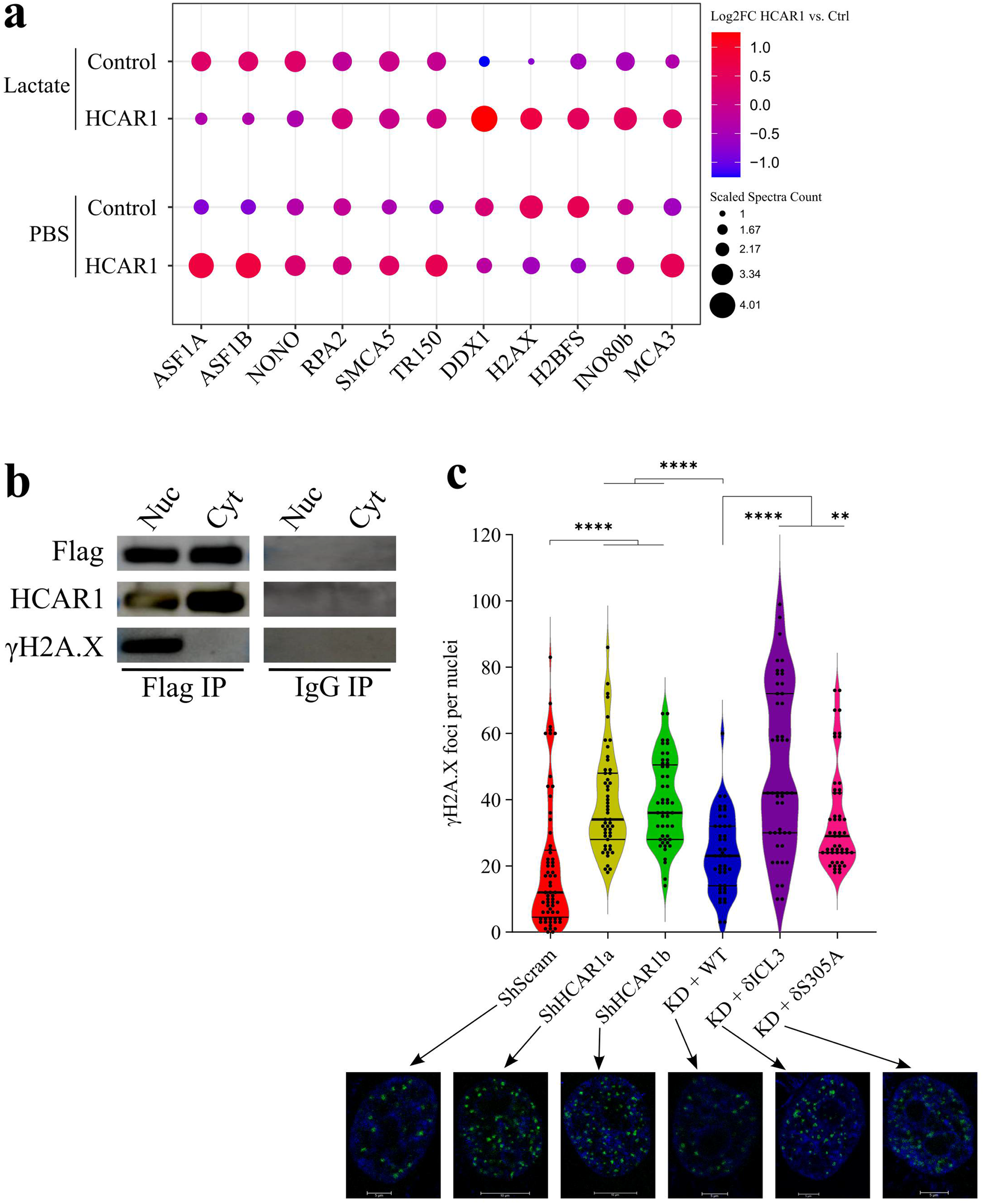
N-HCAR1 with its interactome promotes DNA damage repair. **a)** Dot plot of enriched proteins with HCAR1 which are involved in DNA damage repair. **b)** co-immunoprecipitation of γH2AX with HCAR1 or IgG in fractionated cells. **c)** Irradiated cells were let to recover for 4h and the amount of DNA damage was measured with γH2AX foci. Each dot represents the number of γH2AX foci per nucleus, for 4 separate experiments. Underneath are the representative nuclei of irradiated cells with confocal imaging of γH2AX staining. Data are mean ± s.d. from n=4 biological replicates. ANOVA was followed by Bonferroni post hoc correction test with * *P* < 0.05, ** *P* < 0.01, \*\*\**P* < 0.0001 significance levels.

### N-HCAR1 binds to array of gene complexes, and regulate cell migration

Since several chromatin remodeling factors were detected in the interactome data with the N-HCAR1 (Fig. 3b), we performed genome wide ChIP-sequencing of HCAR1 to identify the genes that are interacting with the receptor. Around 260 genes significantly interacted with HCAR1 upon its stimulation, while the number of genes associated with the unstimulated receptor was much higher (∼600) (Fig. 5a; Supp Fig. 7a-d). Less than 8% of the genes were shared by treatment or not with lactate, inferring that a conformational change in N-HCAR1 caused a genomic redistribution. Consistently, unstimulated N-HCAR1 mostly localizes to gene deserts while upon lactate stimulation it occupies gene segments (Fig. 5b). The same applies within gene regions, where unstimulated N-HCAR1 distributes in an unorganized pattern around transcription start sites compared to a precise reorientation to transcription start sites upon lactate stimulation, suggesting a putative role in gene expression regulation (Fig. 5c). Accordingly, expression analysis for some of the most interactive genes with HCAR1 based on our ChIP-seq analysis upon lactate stimulation reveals an HCAR1-dependent expression profile, abrogated by N-HCAR1 KD (Fig. 5d); this provides additional evidence for direct gene regulatory function of N-HCAR1. Since several epigenetic modifiers were detected in the interactome with N-HCAR1 (Fig. 3b), we questioned and found that the promoter of HCAR1-bound genes are co-enriched with positive regulatory epigenetic markers including H3K9ac, H3K27ac and H3K4me3 (Dunham *et al*., 2012), (Xu, Du and Lau, 2014), but not with compact chromatin marker H3K27me3 (Fig. 5e).

**Fig. 5:**
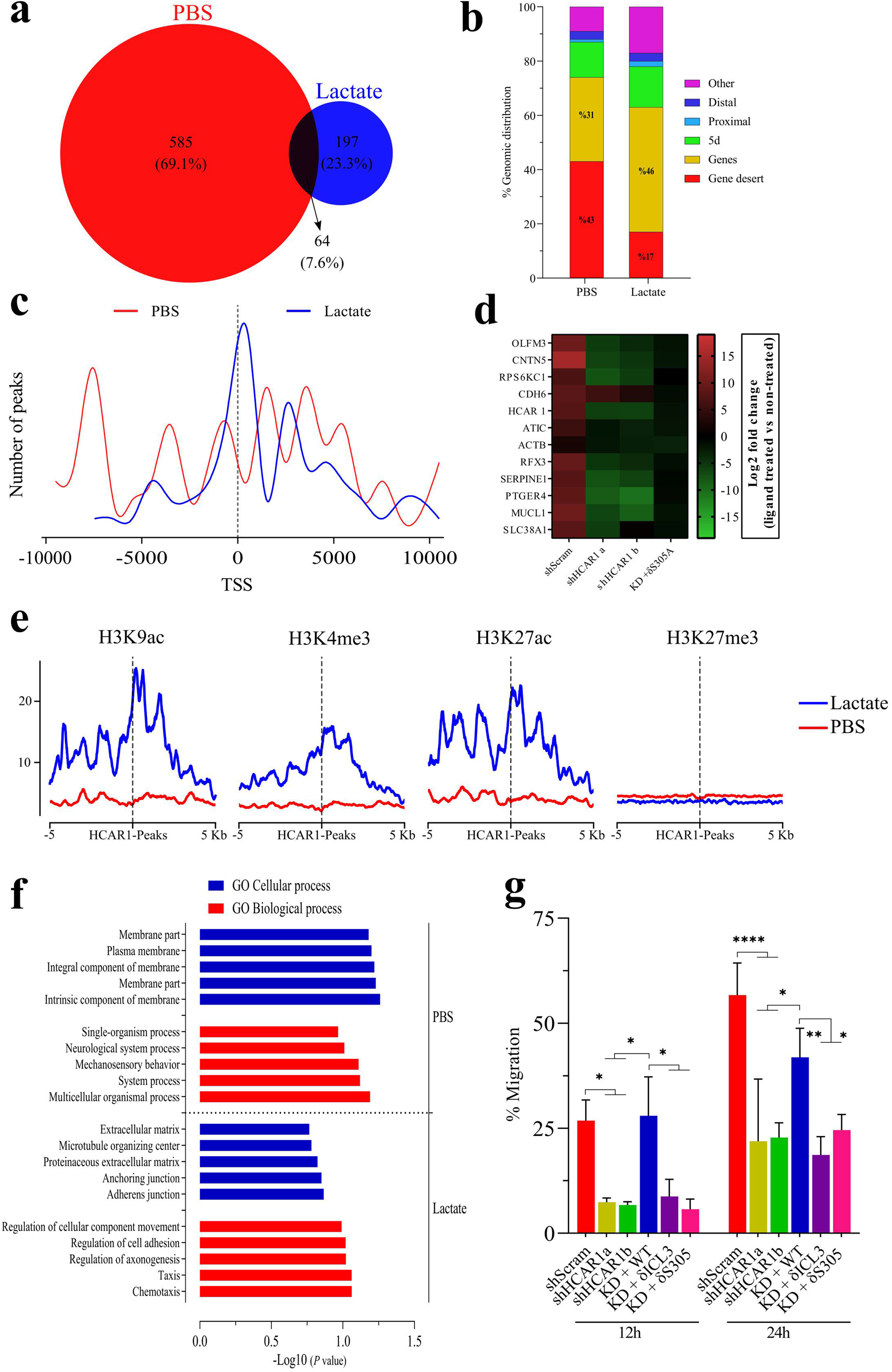
HCAR1 genome-wide interactions show enrichment for genes promoting migration. **a-c)** ChIP-seq of HCAR1 from PBS or Lactate-treated (10mM for 1h) cells from quadruplicate samples. **a)** Venn diagram representing the number of genes associated with HCAR1 in each treatment. **b)** Genomic distribution of HCAR1 in each treatment. Genes (exon or intron), proximal (2kb upstream of TSS), distal (between 2 and 10kb upstream of TSS), 5d (between 10 and 100kb upstream of TSS), Gene desert (≥100kb up or down stream of TSS), Others (anything else). **c)** Normalized number of HCAR1 peaks around TSS of genes. **d)** qRT-PCR for the top 4 genes in each section of the Venn diagram (panel a). Expression levels are presented as Log_2_ fold changes of lactate treated (10mM for 6h) cells over PBS treatment (n=4). **e)** Co-alignment of histone marks from encode project from HeLa cells over HCAR1 peaks. **f)** Ontological analysis of HCAR1-bound genes in PBS- and lactate-treated samples. **g)** Scratch assay to measure the migration rate of cells (n=3). Data in panel d) & g) are mean ± s.d. from biological replicates. Their ANOVA was followed by Bonferroni post hoc correction test with * *P* < 0.05, ** *P* < 0.01, \*\*\**P* < 0.0001 significance levels. TSS: Transcription Start Sites.

Ontological and gene set enrichment analysis reveals that while the enriched genes for unstimulated N-HCAR1 are mainly involved in general homeostatic processes, ligand activated N-HCAR1 binds to genes that regulate various features of cell migration (Fig. 5f) - another component of cell propagation. Cell scratch assay (migration rate test) confirmed this presumption by disclosing limited migration of HeLa cells devoid of total or nuclear HCAR1 (Fig. 5g); similar observations were made on U251MG and A549 cells (Supp Fig. 5e,f). These observations show that N-HCAR1 is able to directly regulate genes involved in cellular movement and promote cell migration.

### N-HCAR1 regulates a larger gene network than its plasma membrane/cytoplasmic counterpart

Our observations for N-HCAR1 mediated intra-nuclear signaling and interactions with various nuclear proteins and genes suggested that N-HCAR1 regulates expression of several genes through various mechanisms. To elucidate the transcriptomic network regulated by N-HCAR1 we performed RNA-seq on cells expressing or not HCAR1 or the N-HCAR1 in presence or absence of lactate (Supp Fig. 7e). Stimulated and unstimulated N-HCAR1 disclosed different transcriptomic profiles (Fig. 6a,b); only ∼34% of genes were shared for stimulated and unstimulated conditions (Fig. 6b). Interestingly of all differentially regulated genes by HCAR1, ∼35% were governed solely by N-HCAR1 and ∼26% through plasma membrane/cytoplasmic HCAR1 (Fig. 6b,c); interestingly, unstimulated N-HCAR1 regulates a larger gene network than its plasma membrane/cytoplasmic counterpart (Fig. 6c). This shows that N-HCAR1 has a more significant function in the nucleus than rest of the cell, at least in terms of transcriptional regulatory output.

**Fig. 6:**
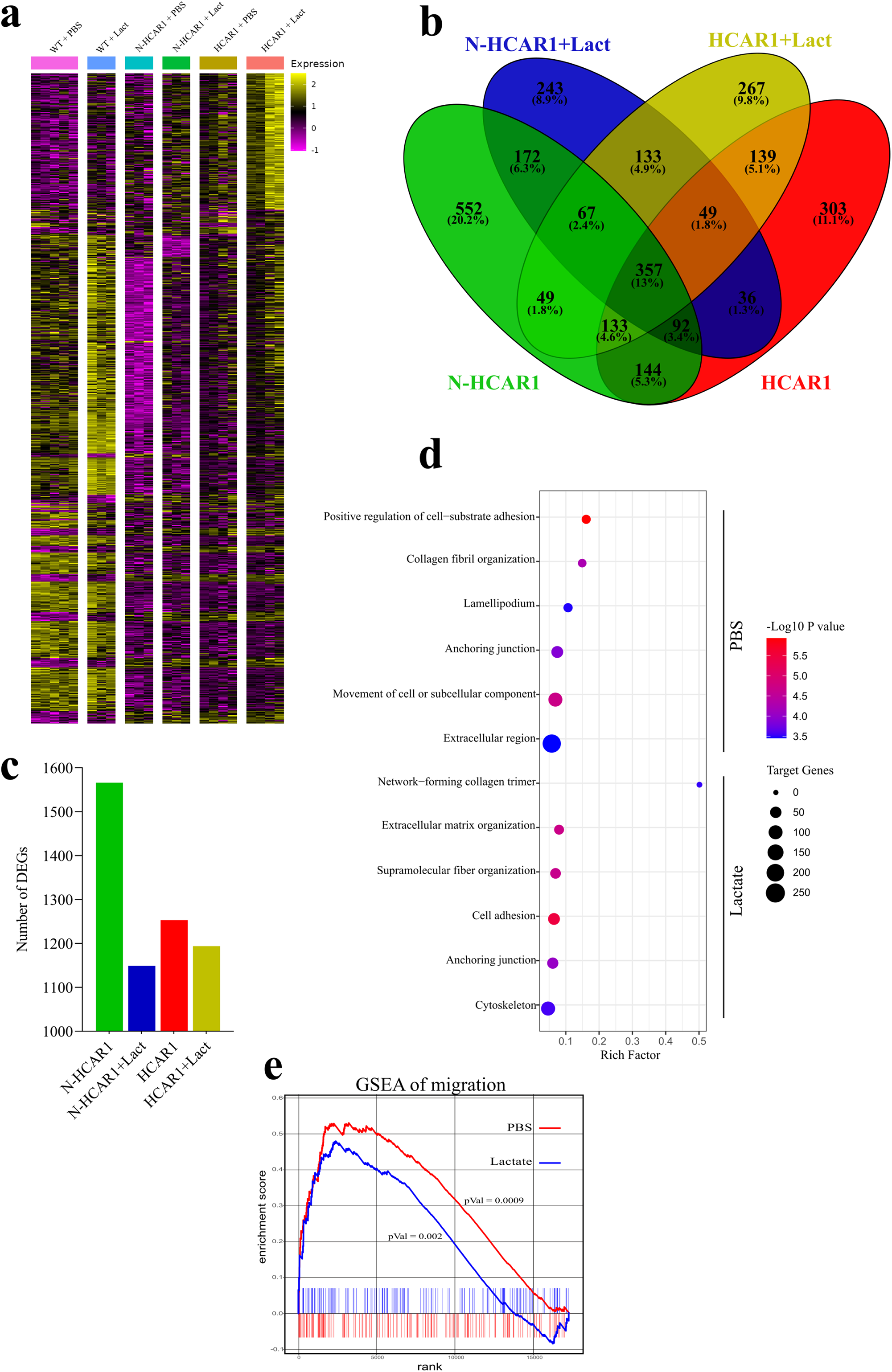
N-HCAR1 regulates a larger gene network than its plasma membrane/cytoplasmic counterpart. **a-c)** RNA-seq of PBS and lactate treated (10mM for 6h) samples from Scrambled shRNA, shHCAR1b and shHCAR1b+ RNAi δS305A HCAR1 cells. **a)** Heatmap of significantly DEGs. **b)** Venn diagram representing all DEGs in each line compared to shScrambled with their corresponding treatment. **c)** Bar graph representing total number of all DEGs in each line compared to shScrambled with their corresponding treatment. **d)** Ontological analysis of genes that were uniquely downregulated only in HCAR1 nuclear KD cells with PBS or lactate treatments. **e)** GSEA waterfall plot of migration. Genes list is extracted from those that are uniquely downregulated only in HCAR1 nuclear KD cells with PBS or lactate treatments. The expression values represent WT condition to indicate expression level of the genes regulated through N-HCAR1. shScrambled PBS (n=5), shScrambled Lactate (n=3), shHCAR1b PBS (n=4), shHCAR1b Lacate (n=4), shHCAR1b+ RNAi δS305A HCAR1 PBS & Lacate (n=3). DEG: Differentially Expressed Genes.

Consistent with ChIP-seq data, ontological analysis of lactate stimulated N-HCAR1-dependent transcriptome related to migration pathways including anchoring junctions, network-forming collagen trimer and extracellular matrix organization (Fig. 6d); transcriptomic analysis of unstimulated N-HCAR1 revealed other aspects of migration such as cell-substrate adhesion, collagen fibril organization and lamellipodium (Fig. 6d). Gene Set Enrichment Analysis (GSEA) on migration ascertained N-HCAR1-dependent (stimulated or not with lactate) upregulation of genes involved in migration (Fig. 6e). Altogether, both stimulated and unstimulated N-HCAR1 coordinately promote expression of genes involved in different features of cellular movement leading to enhanced migration (as per Fig. 5g).

In an attempt to determine if the N-HCAR1-gene complex based on ChIP-seq results culminated in gene expression or suppression, we aligned GSEA of RNA-seq on the ChIP-seq data. Analysis revealed that genes bound to N-HCAR1 (lactate stimulated or not) were mostly upregulated (Supp Fig. 7f,g). Overall, N-HCAR1 upregulates the expression of genes it binds to, indicating it generally acts as a positive regulatory factor for gene expression mostly independent of lactate stimulation.

### N-HCAR1 promotes cancer growth and propagation *in vivo*

HCAR1 has been shown to enhance cancer progression and metastasis *in vivo* (Roland *et al*., 2014),(Lee *et al*., 2016), (Stäubert, Broom and Nordström, 2015); and N-HCAR1 mediates proliferation, survival and migration of cancer cells *in vitro*, as shown above. We validated the *in vivo* role of N-HCAR1 in expressing HeLa cells containing luciferase, by injecting subcutaneously these cells in NOD/SCID/IL2Rγ null (NSG) mice; proliferation and spread of tumors was monitored by bioluminescent live imaging (Fig. 7a). Tumor volume and mass markedly increased in mice injected with HCAR1-expressing WT rescue cells compared to tumors silenced for HCAR1 and N-HCAR1 KD cells (δS305A rescue) (Fig. 7b,c). Coherently, resected tumors expressing HCAR1 at the nucleus exhibited higher proliferation index (Ki-67) and endothelial density (CD31 positivity), and less apoptosis (TUNEL staining), compared to tumors devoid of nuclear HCAR1 (Fig. 7d; Supp Fig. 5). To assess metastatic spread capacity, HeLa cells were injected in the tail vein and metastatic tumor spread was monitored by bioluminescence (Fig. 7e). As seen with tumor volume, metastatic spread was observed only in HeLa cells expressing nuclear-intact HCAR1 and none was detected in HCAR1 KD and N-HCAR1 KD cells. (Fig. 7f). These data support the notion that nucleus-localized HCAR1 exerts a critical role in promoting cancer growth and propagation (malignancy) *in vivo*.

**Fig. 7:**
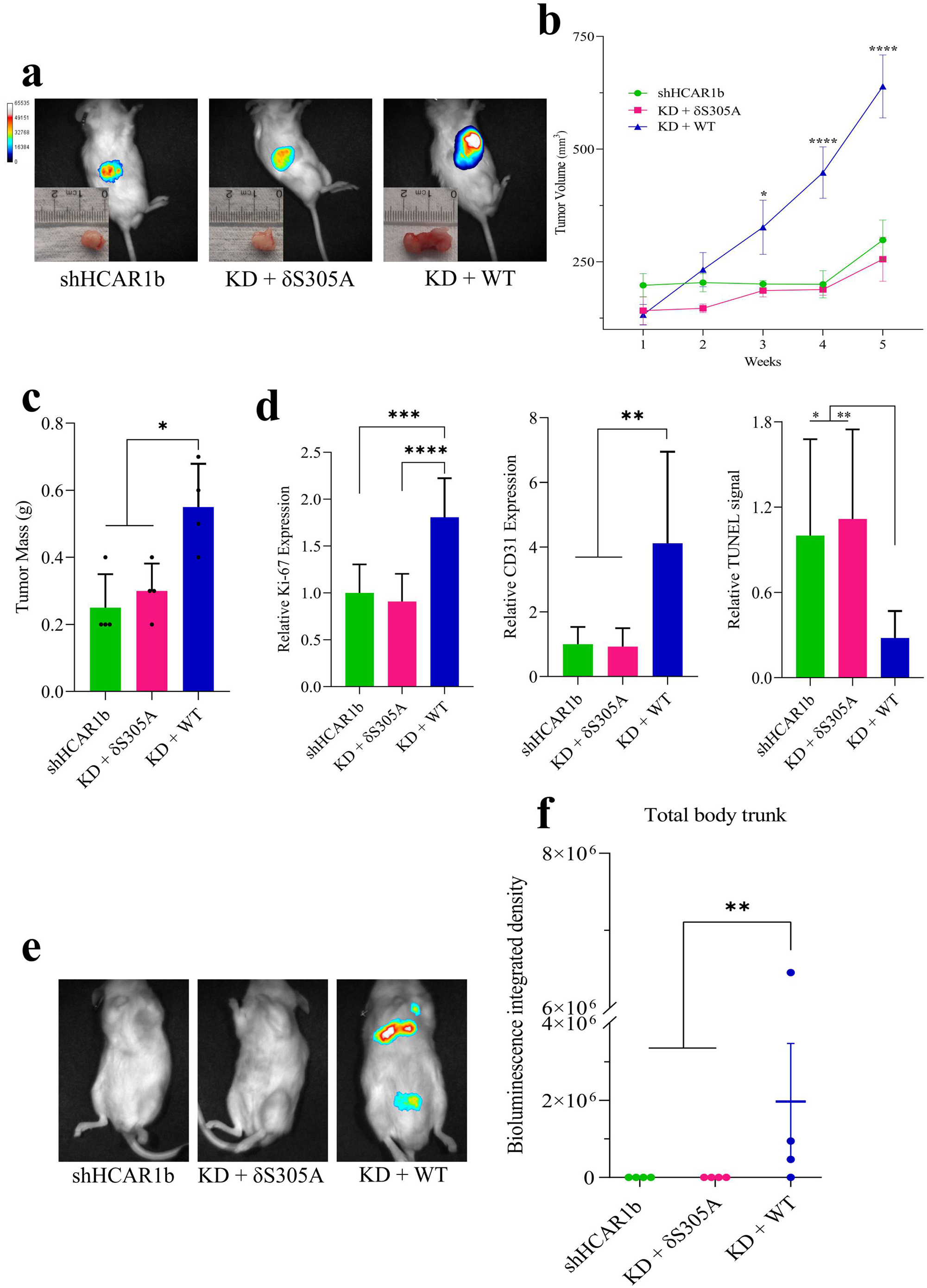
N-HCAR1 promotes cancer malignancy *in vivo*. **a-d)** Subcutaneous injection of luciferase expressing shHCAR1b cells, rescued or not with RNAi-resistant constructs δS305A or WT HCAR1, in NSG mice. **a)** Representative images of *in vivo* luciferase signal and corresponding dissected tumors 5 weeks after injection. **b)** Tumor volume measurement. **c)** Weight of dissected tumors 5 weeks after injection. **d)** Immunohistochemistry staining analysis from dissected tumors indicating relative expression levels of Ki-67 and CD31 and relative cell death (TUNEL assay). **e-f)** Tail vein injection of the same cell lines as above in NSG mice. **e)** Representative luciferase *in vivo* images indicating metastasis of the cells. **f)** Bioluminescence intensity from body trunk of mice indicating metastasis. Each dot represents one mouse. Data in panel b, c & f are mean ± s.e.m. and in panel are mean ± s.d., from n=4 biological replicates. The ANOVA was followed by Bonferroni post hoc correction test with * *P* < 0.05, ** *P* < 0.01, \*\*\**P* < 0.0001 significance levels.

## Discussion

GPCRs are involved in essentially every pathophysiological process; this has largely been based on their plasma membrane location. Mounting evidence points to intracellular location of GPCRs, such that their location-biased activity is arising as an emerging concept (Mohammad Nezhady, Rivera and Chemtob, 2020). This understudied aspect of GPCR biology provides new avenues for therapeutic exploitation of this highly druggable receptor family; location bias expands on the physiologic effects of GPCRs which could explain enigmatic features of their involvement in a number of roles. Counterintuitively, the requirement for intracellular lactate is observed for a number of HCAR1-dependent functions and the receptor’s mechanism of action is unexplained in these conditions (Wagner, Ciszewski and Kania, 2015),(Wagner, Kania and Ciszewski, 2017),(Raychaudhuri *et al*., 2019). Herein, we demonstrate several unprecedented molecular functions for a GPCR at the nucleus, which promote cancer malignancy. While the nucleus contains one third of the cellular reservoir of HCAR1 (Fig. 1e), we provide unmatched evidence that effects of a nuclear GPCR on gene regulation surpasses that through its plasma membrane counterpart, underlining in this case the importance of nuclear HCAR1. Essentially, nuclear HCAR1 beside its signaling activity, directly governs gene regulation for various important functions including migration and through protein-protein interactions modulates critical processes such as protein translation. Consistent with previous reports for other GPCRs (Joyal *et al*., 2014), nuclear HCAR1 regulates genes expression through intranuclear signaling from nuclear membrane as well as its intranuclear sites. But in this case we demonstrate that the GPCR, HCAR1, at the nucleus, also modulates a vast array of cell functions (i.e., protein translation and DNA damage repair) similar to known nuclear receptors such as the estrogen receptor which exert roles in protein translation and splicing in addition to gene regulation (Xu *et al*., 2021).

HCAR1 was found at the INM with analogous conformation to its plasma membrane counterpart and comparably capable of triggering classical G protein-coupled signaling bursts of ERK and AKT activation in the nucleus. Furthermore, intranuclear HCAR1 interacted with a variety of proteins involved in various physiologic functions such as ribosomal biogenesis, protein translation rate, DNA damage repair and with genome-wide chromatin-binding complexes involved in cell migration. Based on interactome data, HCAR1 binds to different transcriptional factors from INO80, SWI/SNF and ISWI ATP-dependent chromatin remodeling complexes such as INO80b, SMARCA5, SMARCC1 and BPTF (Hasan and Ahuja, 2019); these interactions were observed with the activated and non-activated receptor, suggesting a potential constitutive role of HCAR1 in modulating their activity in a metabolic-dependent manner. While these chromatin remodelers are significantly miss-regulated in many cancers (Lu and Roberts, 2013),(Prendergast *et al*., 2020),(Li *et al*., 2021), N-HCAR1 activated by higher concentration of lactate seen in tumor (Warburg effect) could alter their activity (in favor of cancer promotion). Interestingly, inactive N-HCAR1 also interacted with NSD1, a histone methyltransferase known to bind to different nuclear receptors (including estrogen, thyroid, retinoic acid, and retinoid receptors) (Tauchmann and Schwaller, 2021); since NSD1 is frequently mis-regulated in cancers (Tauchmann and Schwaller, 2021) it is tempting to speculate that metabolic rewiring could cause epigenomic alterations in favor of cancer malignancy. Concordantly, our genome-wide association study shows how N-HCAR1 could directly promote expression of genes involved in migration potentially through these epigenetic modulations. We found that several genes such as *WNT3* (Madaan *et al*., 2019),(Lee *et al*., 2018), *SERPINE1* (Lee *et al*., 2016), *CDH5* (Khatib-Massalha *et al*., 2020) previously reported to be regulated through HCAR1, are bound by N-HCAR1 suggestive of direct gene regulation. Direct gene regulation also seems to apply to *HCAR1* as well through lactate-stimulated N-HCAR1, consistent with reported auto-induction of HCAR1 (Zhao *et al*., 2020),(Liu *et al*., 2009), (Ishihara *et al*., 2022).

GPCRs with ligands constantly present, are stochastically in both active and inactive states at any given time; the ratio of active to inactive state depends on the concentration of ligand (Felce, Davis and Klenerman, 2018). On the other hand, a single molecule GPCR stoichiometry can simultaneously bind Gα, Gβ, GRKs, and arrestin (Gurevich and Gurevich, 2018). These interactions occur through the intracellular domains of a GPCR, and are distinct from their ability to form homo/heterodimers via their hydrophobic transmembrane domains (Wu *et al*., 2010), thus providing other docking sites for protein-protein interactions. These intricacies expose the abilities of GPCRs to form various protein complexes. Along these lines, one could envisage non-cylindrical conformations for a GPCR inside the nucleus with its hydrophobic domains deeply buried in protein complexes interacting with hydrophobic domains of other proteins (Mohammad Nezhady, Rivera and Chemtob, 2020); such interactions could explain N-HCAR1’s transcriptional and translational control from within the nucleus.

The present study highlights the multifaceted functionality of GPCRs through its nuclear location; nuclear HCAR1 provides an adaptive fitness of cells to respond to metabolic tweaks through intracellular ligands, as is the case for lactate which augments survival, proliferation and propagation of cancer cells, by acting via N-HCAR1. These myriad of roles for N-HCAR1 might not be determinant for individual cellular processes it participates in, however its collective functions on various processes convey a significant adaptation for cancer progression and malignancy.

## Acknowledgment

This work was supported by Canadian Institutes of Health Research grant. MA.MN was supported by S.Véronneau□Troutman & Université de Montréal Ophthalmology department, program in molecular biology and faculty of medicine scholarships. M.M was supported by scholarships from faculty of medicine Université de Montréal. U251MG cells was a gift from Dr. Hardy’s lab. We thank Dr. E. Kuster from microscopy facility of CR Sainte Justine Hospital, Dr. R. Lambert from genomic facility at IRIC, Dr. E. Bonneil from IRIC proteomics, & Dr. D. Gingras from Electron microscopy facility at UdeM and Dr. X. Hou for their inputs on the work presented here. We thank D. Obari from i-science.ca for the graphic work. SC holds a Canada Research Chair (Translational Research in Vision) and the Leopoldine Wolfe Chair in translational research in age-related macular degeneration.

## Contribution

MA.MN conceived and designed the study, performed and analyzed the experiments and drafted the manuscript. G.C performed the bioinformatic analysis under supervision of S.J. E.B & P.C contributed to the analysis of the data. M.M helped with performing animal study. D. M provided the Luciferase virus and A549 cells and helped with the analysis of some data under the supervision of C.B. S.C supervised the whole project and oversaw the conception, experiments, analysis and drafting of the work.

## Figure Titles and legends

**Supp Fig. 1:**

**a-b)** Control experiments for HCAR1 localization with empty vector containing Flag tag (a), and no primary antibody staining control (b) in immunofluorescence confocal imaging. **c-d)** Immunofluorescence confocal imaging of N-terminally Flag tagged HCAR1 in isolated nuclei (c) and whole cell (d). **e-f)** Control experiments for HCAR1 localization with empty vector containing Flag tag (e), and no primary antibody staining control (f) in TEM images. g-h) TEM images of N-terminally Flag-tagged HCAR1 with PBS (g) and Lactate treated (10mM for 1h) cells (h). All images are in HeLa cells.

**Supp Fig. 2:**

**a-c)** Immunofluorescence confocal imaging of C-terminally Flag tagged HCAR1 in U251MG cells with WT HCAR1 (a), δICL3 HCAR1(b) and δS305A HCAR1 (c) cells. **d-f)** Immunofluorescence confocal imaging of C-terminally Flag tagged HCAR1 in A549 cells with WT HCAR1 (d), δICL3 HCAR1 (e) and δS305A HCAR1 (f) cells.

**Supp Fig. 3:**

**a-d)** Confocal imaging of pulse-chase FAP system with HCAR1 in HeLa cells using impermeant green fluorogen followed by lactate treatment for 5 to 40min. **e)** Immunofluorescence confocal imaging pMFAP-β1 FAP construct with C-terminally Myc-tagged HCAR1 in HeLa cells.

**Supp Fig. 4:**

**a)** Immunofluorescence confocal imaging of isolated nuclei with selective ONM permeabilization. Detection of Sun2 C-terminus indicates INM permeabilization and the absence of Lamin B1 signal indicates intact non-permeabilized INM. **b)** Western-blot analysis on isolated nuclei from WT cells, cells overexpressing C- & N-terminally tagged HCAR1, shScrambled or two HCAR1 KD HeLa cells. Isolated nuclei were stimulated with PBS or lactate (10mM for 15min). **c-d)** Western-blot analysis on isolated nuclei from shScrambled HeLa cells from different treatments. **e)** Schematic of the modifications in the HeLa cell lines used in this study. WT HCAR1-expressing HeLa cells were stably transduced with shRNA against 3’UTR of HCAR1 (and scrambled control of shRNA) to stably KD the HCAR1. Then, these cells were transfected with RNAi resistant plasmid expressing either the coding sequence of WT or the mutant versions of the HCAR1, to generate stably expressing cells that either harbor WT or nuclear excluded HCAR1. **f)** qRT-PCR analysis of *HCAR1* expression normalized to WT validating KD and rescue efficiencies. **g)** Quantitation western blot analysis of HCAR1 expression validation KD and rescue efficiencies. **h)** *HCAR1* expression level in 3 cell lines used in this study; nTPM: normalized protein-coding transcripts per million. (proteinatlas.org)

**Supp Fig. 5:**

**a-b)** Homeostatic proliferation rate (left panel) and survival rate in 5FU-treated cells (right panel) in U251MG (a) and A549 (b) cell lines. Both U251MG and A549 cell lines are expressing lower levels of endogenous *HCAR1* (see Supp Fig. 4h), so we generated stable cell lines over-expressing either WT HCAR1, or nuclear-excluded δICL3 HCAR1 and δS305A HCAR1 in these two cell lines. **(c-d)** Protein translation rate with methionine incorporation rate measurement in U251MG (c) and A549 (d) cell lines. Methionine incorporation rate (AHA) was adjusted to the number of cells (Hoechst). **(e-f)** Scratch assay to measure the migration rate in U251MG (e) and A549 (f) cell lines. Migration rate was measured 8h post-scratch in U251MG cell lines and 18h post-scratch in A549 cell lines.

**Supp Fig. 6:**

**a)** Immunofluorescence confocal imaging of HA-tagged HCAR1-BirA construct shows same localization pattern (on the nuclear membrane and in inside the nucleus) for HCAR1-BirA fusion protein as the WT HCAR1 in HeLa cells. **b)** Western blot analysis with streptavidin-HRP on biotinylated whole cell lysate from PBS or Biotin-treated cells. **c-d)** Heatmaps showing enrichment of proteins with HCAR1 based on Log_2_ Fold change and *p* value in isolated nuclei of PBS (c) or Lactate-treated (d) cells with biotin. Control samples are from stable cell lines expressing empty Bio-ID vector, expressing only BirA. **e)** Enrichment dot plot graph showing proteins enriched in tRNA aminoacylation pathway in PBS and lactate-treated cells compared to control cells. **f)** Enrichment dot plot graph showing proteins enriched in ribosome biogenesis pathway in PBS and lactate treated cells compared to control cells.

**Supp Fig. 7:**

**a-c)** ChIP-qPCR confirmation for ChIP-seq. Selected genes are the top 4 enriched loci in each PBS-treated only (a), Lactate-treated only (b), and shared genes (c). **d)** Integrative Genomic Viewer (IGV) of the *HCAR1* locus in the ChIP-seq data. Peaks shows HCAR1 is bound to its gene both in PBS- and Lactate-treated cells, but there is no enrichment in input samples. **e)** Expression level of *HCAR1* in RNA-seq data validating the RNA-seq as it shows its low expression in KD (bL: shHCAR1b+ Lactate; bP: shHCAR1b+PBS), but higher expression in scrambled shRNA and rescue of KD with δS305A HCAR1 (dsL, dsP). **(f-g)** Waterfall plots representing overall general positive regulatory function of N-HCAR1 on gene transcription in N-HCAR1 bound genes. The expression values are extracted from RNA-seq data of HCAR1 nuclear KD cells with PBS and lactate treatments. The expression values represent WT condition to indicate expression level of genes regulated through N-HCAR1. The gene list is extracted from PBS-treated (f) and Lactate-treated (g) HCAR1 ChIP-seq data.

## Material and Methods

### Cell lines and treatments

HeLa (CCL-2 ATCC) and A549 (CCL-185 ATCC) were purchased from commercial vendors and maintained according to the manufacturers protocol in DMEM, 10% FBS and 1% Pen/Strep, U-251MG cells were a kind gift from Dr. Hardy’s lab and maintained in EMEM + 2 mM Glutamine + 1% non-essential amino acids + 1 mM Sodium Pyruvate + 10% FBS and 1% Pen/Strep in a humidified incubator with 5% CO2 at 37□°C. Stable cells were generated using appropriate drug selection (G-418, Puromycine) after plasmid transfection or viral transduction, and were maintained in these antibiotic instead of Pen/Strep. A stock concentration of 500 mM lactate (in PBS and pH adjusted to 7.4) was used for cell stimulation and similar volume of PBS as vehicle was used as control. The data for end point phenotypic effects in HeLa cell are presented in the main figures and the data for end point phenotypic effects in A549 and U-251MG cells are presented in the extended figures.

Cell replication and survival were determined by enumerating live and dead cells using automatic countess cell counter (Thermofisher) using trypan blue exclusion assay (Thermofisher). Cells were treated with 20μM 5FU or starved for 24h for survival assay, and cell numbers were calculated before and after the treatments. Trypan blue was added in 1X ratio to the media containing cells and each replicate was performed in quadruplicates to determine the number of live and dead cells. Each experiment was conducted at least in triplicates.

### Plasmids, RNAi and CRISPR

Cells were transfected with human HCAR1 because of enhanced immunoreactivity to exogenous tag (such as Flag), enabling superior localization resolution and for immunoprecipitation, as well as for FAP and Bio-ID construct preparation. The cDNA encoding HCAR1 was PCR-amplified with encompassing appropriate restriction enzymes sites at both ends of the amplicon. The final product was gel-purified (ThermoFisher GeneJET Gel Extraction Kit), digested with the restriction enzymes and cloned into each vector. pCMV-Tag 2A (Agilent) was used for N-terminal flag tagging using EcoRI and HindIII flanking sites, pCDNA3.1-HCAR1-flag (Genscript) was used for C-terminal flag tagging. ICL3 and S305A mutations were generated using back-to-back primers on pCDNA3.1-HCAR1-Flag vector by Q5 Site-Directed Mutagenesis Kit (NEB). Fluorogen activating peptide fusion to HCAR1 was synthesized with insertion of HCAR1 using BsmI site into pMFAP-β1 vector (Spectragenetics), and BioID fusion was generated with HCAR1 insertion into flanking sites AccIII and AfIII in MCS-13X Linker-BioID2-HA (Addgene 80899) vector. All plasmids were sequenced to verify the correct insertion. Vectors were transfected into the cells using TranIT-X2 reagent (Mirus) according to the manufacturers protocol and grown on appropriate antibiotics to generate stable cell lines.

Lentiviral shRNA against HCAR1 targeting 3’UTR regions of the gene (shHCAR1a: GCTTTATTTCAGGCCGAATGA; shHCAR1b: GCTCTGACCTTCTTCAAATCT) and the scrambled shRNA were purchased from GeneCopoeia (Cat# LPP-HSH007585-LVRU6MP-100). Targeting the 3’UTR regions allowed us to use our previous plasmid constructs for rescue experiments.

### RNA isolation and quantitative PCR

RNA was isolated using either RiboZol (VWR) or RNeasy mini kit (Qiagen) then was converted to cDNA using iScript (Bio-Rad) following manufacturer’s instructions. qRT-PCR was performed using SYBR green master mix (Bio-Rad) on Roche light cycler. HRP and 18S were used for normalization of the results (normalization to 18S is reported in the manuscript).

### Immunoblot and ELISA

Cells were lysed in RIPA buffer (Cell Signaling) and a cocktail of protease inhibitors (Roche) and protein concentration was measured with Bradford assay (Bio-Rad). Proteins were heated in reducing Laemmli sample buffer at 95°C and resolved in SDS-PAGE protein gel and transferred to PVDF membrane (Bio-Rad). Membranes were blocked using 5% BSA (Sigma) for 1 h and then incubated for overnight with the primary antibodies. Afterward membranes were washed 3X with TBST and incubated with HRP-conjugated secondary antibodies for 1h and then were washed again and revealed by ECL (VWR) chemiluminescence.

ERK1/2 and AKT phosphorylation levels were measured with both western blot and ELISA kits (Abcam). Cells were treated overnight with PTX (300ng/ml) or Gallein (20μM) and then nuclei were isolated and suspended in 10mM lactate or vehicle with rotation at 37°C for 15min, washed with PBS 2X and then were lysed in either Laemmeli buffer for western blot or treated according to the manufacturers protocol for ELISA.

### Immunofluorescence (IF) staining

Cells were seeded on Poly-L-Lysine coated cover slips in a 6 well plates for over night in incubator. For DNA damage, cells were irradiated with 1Gy intensity using Faxitron CP-160 irradiator and let to recover for 4 hours at 37°C in incubator, and then IF was performed on them. Wells were rinsed three times with PBS, fixed in 4% formaldehyde (Sigma) for 10 min at RT, and then washed three times 5min with PBS. Subsequently, cells were permeabilized with 0.1% Triton X-100 in PBS for 15 min at RT and blocked in 1% BSA in PBS/ 0.1% Tween-20 for 1h. Cover slips were incubated in a humid chamber with primary antibody for overnight at 4°C diluted in new blocking buffer. The antibody solution was washed with PBST (3x, 5min) and samples were incubated with secondary antibody (Alexa-fluor conjugated secondaries) for 1h at RT in dark. Cells were washed again, stained with DAPI, washed and then mounted. Slides were imaged with Leica confocal microscopy (SP8 or SP8-STED) with appropriate channels and 60X objectives. Fluorescent lights were gated to avoid any overlap between channels. Images were analysed using LAS-X (Leica microscopy licenced software) and ImageJ software.

### TEM

Cells were fixed in 4% paraformaldehyde + 0.5% Glutaraldehyde in cacodylate buffer (0.1M, pH7.2). After fixation, cells were washed 2 times in cacodylate buffer (5 min) and then in PBS. Permeabilization was performed with 0.2% Triton X-100 for 15 min, and then cells were blocked with PBST with 10%FBS for 1h. Samples were incubated ON with primary antibody in blocking buffer at 4°C, and delivered to electron microscopy facility of faculty of medicine (Universite de Montreal) for further processing and imaging. Primary antibody was detected with nanogold conjugated secondaries and silver enhancement, and imaged with transmission electron microscope (Philips CM120) equipped with a Gatan digital camera.

### Fluorogen Activating Peptide (FAP) pulse chase

HCAR1 gene was cloned into pMFAP-β1 vector (Spectragenetics) and pulse chase experiments were performed after generation of stable cell lines. Cell were treated with 100nM βGREEN-np membrane impermeant fluorogen (Spectragenetics), which can not enter the cell unless bound to FAP, and one minute later cells were treated with 10 mM lactate (or PBS) for indicated time points. Afterward, cells were briefly washed with PBS and fixed with 4% paraformaldehyde, and the nuclei was stained with Hoechst 33342. βGREEN-np was excited and imaged using Alexa fluor 514 channel with Leica confocal microscopy. No nuclei with βGREEN-np were observed in the experiments. IF staining of stable cell lines without lactate treatment were performed as mentioned previously with Myc primary antibody against the Myc-tag in the N-terminus of HCAR1 and C-terminus of FAP.

### Nuclei isolation & staining

Isolating nuclei was performed as described previously(Nabbi and Riabowol, 2015). Briefly, cells were washed and resuspended in PBS plus protease inhibitor cocktail (Roche) and 1mM PMSF. They were centrifuged (10000rpm for 10sec) and resuspended in PBS + protease inhibitor cocktail + 0.1% NP-40 and triturated for 7 times with P1000 micropipette tip. Supernatant was collected (or removed) after centrifugation as cytoplasmic fraction. The pellet (containing nuclei) was subjected to second time trituration (5 times) and centrifuged again to obtain pure nuclei fraction. After last centrifugation, supernatant was removed and nuclei was collected in the desired solution. The purity of nuclear fraction was checked under the microscope and was validate by western blotting.

Intact non-permeabilized nuclei were resuspended in PBS and mounted on Poly-L Lysine coated cover slips for IF staining. For ONM permeabilization, nuclei were resuspended in 0.0008% Digitonin and rotated for 5min at RT in microfuge tubes. Permeabilized nuclei were centrifuged, washed and resuspended in PBS + PI cocktail. For proteinase K digestion, nuclei were incubated at 37°C with rotation in 100μg/ml proteinase K (Sigma) solution before or after ONM permeabilization with Digitonin. Nuclei were then washed 3x in 1%BSA + 5mM PMSF solution and then further permeabilized with Digitonin for ONM permeabilization or 0.1% triton for INM permeabilization and then were subjected to IF staining.

### HCAR1 3D modeling

We analyzed the structure of HCAR1 in both active and inactive forms as described elsewhere(Miszta *et al*., 2018),(Kuei *et al*., 2011), using the web service: https://gpcrm.biomodellab.eu/. Also we examined the inactive structure using recent AlphaFold(Jumper *et al*., 2021), and there was minimal differences between both models only in low confidence regions of the free c-terminus. Post-translation phosphorylation site analysis was done using: https://www.phosphosite.org/, and described in (Hornbeck *et al*., 2015). Only phosphorylation residues with validated mass spectrometry data (HTP) were used for mutagenesis analysis.

### cAMP measurement in isolated nuclei

Isolated nuclei were resuspended in PBS with 10mM lactate with rotation at 37°C for 10min, nuclei were then counted using hemocytometer and were subjected to immunoassay cAMP Direct kit (Abcam) according to manufacturer’s protocol. Protein G-coated plated were used for the ELISA provided in the kit and measurements were performed with HRP development by measuring its OD at 450nm with Clariostar plate reader.

### Bio-ID & Mass Spectrometry

HCAR1 gene was cloned into MCS-13X Linker-BioID2-HA (Addgene 80899) vector. After validating the fusion protein has same localization pattern as the HCAR1 itself, stable cells were generated by antibiotic selection. Control samples were transfected stable cell lines with the empty vector. Cells were treated with Biotin (50μM) and 10mM lactate (or PBS) and incubated for ∼16 hours in incubator. Nuclei were isolated and their purity was validated. Isolated nuclei were lysed with non-denaturing lysis buffer (20 mM Tris HCl pH 8, 137 mM NaCl, 1% Nonidet P-40 (NP-40), 2 mM EDTA) plus PI cocktail. The lysate was incubated with magnetic streptavidin MyOne Dynabeads (ThermoFisher) at 4°C for ON with rotation. Afterward, samples were washed 6x with PBST and then eluted. Eluted samples were sent to LC-MS/MS at IRIC Center for Advanced Proteomics Analyses, a Node of the Canadian Genomic Innovation Network that is supported by the Canadian Government through Genome Canada. Both experimental and control samples were analyzed in triplicates.

BioID eluted samples were reconstituted in 50 mM ammonium bicarbonate with 10 mM TCEP [Tris(2-carboxyethyl) phosphine hydrochloride; Thermo Fisher Scientific], and vortexed for 1 h at 37°C. Chloroacetamide (Sigma-Aldrich) was added for alkylation to a final concentration of 55 mM. Samples were vortexed for another hour at 37°C. One microgram of trypsin was added, and digestion was performed for 8 h at 37°C. Samples were dried down and solubilized in 4% formic acid (FA). Peptides were loaded and separated on a home-made reversed-phase column (150-μm i.d. by 200 mm) with a 56-min gradient from 10 to 30% ACN-0.2% FA and a 600-nl/min flow rate on an Easy nLC-1000 connected to an Orbitrap Fusion (Thermo Fisher Scientific, San Jose, CA). Each full MS spectrum acquired at a resolution of 60,000 was followed by tandem-MS (MS-MS) spectra acquisition on the most abundant multiply charged precursor ions for a maximum of 3s. Tandem-MS experiments were performed using collision-induced dissociation (CID) at a collision energy of 30%. The data were processed using PEAKS X (Bioinformatics Solutions, Waterloo, ON) and the Uniprot human database (20349 entries). Mass tolerances on precursor and fragment ions were 10 ppm and 0.3 Da, respectively. Fixed modification was carbamidomethyl (C). The data were visualized with Scaffold 4.0 (protein threshold, 99%, with at least 2 peptides identified and a false-discovery rate [FDR] of 1% for peptides).

BioID data were analyzed first with Scaffold (Proteome Software Inc. Portland OR) to produce quantitative values from normalized total spectra (Top 3 area based on total ion chromatogram) for the amino acid sequences detected by mass spectrometry. Quantitative values for annotated proteins were then batch corrected using an empirical bayes framework (ComBat, SVA, https://rdocumentation.org/packages/sva/versions/3.20.0) and differential protein abundance between condition was assessed using MetaboAnalyst after quantile normalization, log transformation and autoscaling(Pang *et al*., 2021). Statistical analysis using t-test was performed to determine the proteins enriched in HCAR1 BioID cells compared to cells containing empty vectors (i.e., only the biotin ligase), in PBS and lactate treatment separately. Visualization of normalized data was done in R (version 4.1.0, 2021 The R Foundation for Statistical Computing) using gplots/heatmap.2 (https://CRAN.R-project.org/package=gplots), ggplot2 (https://CRAN.R-project.org/package=ggplot2) and EnhancedVolcano (Blighe K, Rana S, Lewis M (2022). EnhancedVolcano: Publication-ready volcano plots with enhanced colouring and labeling. R package version 1.14.0). Pathway analysis of enriched proteins was performed on EnrichR(Xie *et al*., 2021).

### Ribosomal profiling

Ribosome profiling was performed by sucrose gradient fractionation as described previously (Panda, Martindale and Gorospe, 2017). Briefly, cells were treated with 10μg/ml of cycloheximide (CHX) for 15min at 37°C in the incubator to install ribosome disassembly. Cells were washed and resuspended in cold PBS containing CHX and PI cocktail, and then lysed in lysis buffer (20 mM Tris-HCl,100 mM KCl, 5 mM MgCl2, 0.5% Nonidet P-40) containing CHX, RNase inhibitor and PI cocktail. Lysate was cleared by centrifugation and equal amounts were layered on top of a cold sucrose gradient (10 to 60 % gradient containing CHX, RNase inhibitor and PI cocktail). Gradients were centrifuged in Hitachi swinging ultracentrifuge (CP90NX) at 190,000g for 1.5h at 4°C. Gradients were fractionated by piercing the bottom of sucrose gradient tube and the OD of collected fractions were measured at 254nm spectrum. Ribosomal profile was plotted and area under the curve of each monosome subunit (40S, 60S and 80S) and polysomes were measured for quantification of the ribosomal content.

### Protein translation rate measurement

Nascent protein synthesis rate was measured using Click-iT AHA Alexa Flour 488 protein synthesis HCS assay kit (Invitrogen) according to the manufacturer’s protocol. Equal number of cells were plated ON in a 96-well plate and the media was washed out the next day and replaced with a methionine-free media containing L-azidohomoalanine (AHA) as the methionine analog, and incubated for 30min. AHA is incorporated into proteins during protein synthesis in the methionine-free media. The amount of incorporated AHA is detected with a click chemical reaction by Alexa flour 488. The intensity of Alexa fluor 488 is adjusted with the intensity of DNA counterstain Hoechst 33342 and directly corresponds to the nascent protein synthesis rate.

### Co-IP

Cells were fractionated and cytoplasmic and nuclear fractions were lysed with non-denaturing lysis buffer plus PI cocktail. The lysates were then pre-cleared with equilibrated protein G magnetic beads (Cell Signaling) for 1h at RT with rotation. The pre-cleared lysate was incubated with primary antibody O/N at 4°C. Pre-washed magnetic beads were added to the immunocomplexes and incubated for 1h at RT with rotation. Afterward, beads were isolated with magnetic separation rack and washed 5x with lysis buffer. Finally, beads were resuspended in 3x SDS sample buffer and incubated at 95°C for 5min to elute the immunocomplexes. Elutes were analyzed by western blotting.

### ChIP-Seq

Chromatin immunoprecipitation (ChIP) was performed using SimpleChIP® Enzymatic Chromatin IP Kit (Cell Signaling) based on manufacturer’s protocol. Briefly, DNA-protein complexes were crosslinked using 1% final concentration of formaldehyde (Sigma) for 10min at RT, and quenched with Glycine (final concentration of 125mM). Cells were then washed with cold PBS + PI cocktail, scraped into conical tubes and centrifuged to remove the supernatant. After isolation, nuclei were treated with micrococcal nuclease to digest the DNA, and then sonicated. Digested chromatin was analyzed by agarose gel. Chromatins were then incubated with immunoprecipitating antibody O/N at 4°C with rotation. Control samples were incubated with IgG antibody. ChIP-grade protein G magnetic beads were added to the IP reactions and incubated for 2h at 4°C with rotation. Beads were then washed with low- to high-salt wash buffers and chromatin was eluted in elution buffer for 30min at 65°C. Chromatins were reverse-crosslinked with NaCl and proteinase K and incubation at 65°C for 2h. DNA was purified with spin columns, and analyzed by qPCR or sent to Next Generation Sequencing. Samples were analyzed with bioanalyzer for quality control and single-end NGS was performed at IRIC genomic platform with Nextseq 500 illumina system. Samples were sequenced with a depth of ∼35M per sample with 75 cycles. Both C- and N-terminus flag tagged cells were used for ChIP-seq studies. Two samples from each terminus tagged cells were used for either lactate or vehicle (PBS); in total 4 samples per treatment group, and only shared genes in each group was used for further bioinformatic analysis.

ChipSeq fastq files were processed with default parameter using the ChipSeq pipeline from GenPipe(Bourgey *et al*., 2019). Bam files were visualized with IGV(Robinson *et al*., 2011) (A public access version is also available: PMC3346182). Peak files were analyzed using the ComputeMatrix function from deeptool(Ramírez *et al*., 2016) to determine distance relative to histone modification marks based on Broad Histone Helas Chip-seq data from the Encode project(Luo *et al*., 2020).

### Migration assay

Cells were cultured to reach a density of 90-100% confluency and then a wound was made by scratching the monolayer cells with sterile p200 pipette tips. Cells were washed and new media was added. Cells were imaged by phase contract microscopy right after scratch to measure the initial distance (t0), and later in indicated time points. Reduction of the scratched area due to the migration of the cells were measured as the rate of migration. Each experiment was conducted in triplicates and each time in multiple wells of the plates.

### Transcriptomic

Equal number of cells were seeded in 10cm dish and let to grow a density of 70-80% confluency. Lactate was added to the final concentration of 10mM (or equal volume of PBS) and incubated to 6 hours. RNA was extracted with RNeasy mini Kit (Qiagen). Samples were sent to IRIC genomic platform for analysis and sequencing. RNA integrity and quantity was validated with Bioanalyzer and then used for sequencing. Samples were sequenced with Nextseq 500 illumina system with a depth of ∼35M per sample with single-end 75 cycles. Each sample was sequenced at least in triplicates.

RNAseq data were pre-processed using the RNAseq next-flow pipeline(Ewels *et al*., 2020) with the star_salmon aligner and the salmon pseudo-aligner (reference genome GRCh38). Gene counts were normalized and scaled to perform differential gene expression analysis between groups using Seurat after regressing for batch effect(Hao *et al*., 2021). Differentially expressed genes were further analyzed using fastGSEA(Korotkevich *et al*., 2021) and enrichR(Xie *et al*., 2021).

### Sequencing

500 ng of total DNA for ChIP-sequencing or RNA was used for library preparation. DNA/RNA quality control was assessed with the Bioanalyzer Nano assay on the 2100 Bioanalyzer system (Agilent technologies) and all samples had a RIN above 9,5. For RNA, PolyA selection was done using Dyna Beads Oligo(dT) (Thermo Fisher). Library preparation was done with the KAPA DNA or RNA Hyperprep kit (Roche). Ligation was made with Illumina dual-index UMI (IDT). All libraries were diluted and normalized by qPCR using the KAPA library quantification kit (KAPA; Cat no. KK4973). Libraries were pooled to equimolar concentration. Sequencing was performed with the Illumina Nextseq500 using the Nextseq High Output 75 (1×75bp) cycles kit. Around 30M single-end PF reads were generated per sample for ChIP-sequencing and around 35M for RNA sequencing. Library preparation and sequencing was made at the Institute for Research in Immunology and Cancer’s Genomics Platform (IRIC).

Raw base calls were converted to FASTQ files using bcl2fastq version 2.20 and allowing 0 mismatches in the multiplexing barcode. Prior to that, base calls had been obtained from the Illumina NextSeq 500 sequencer that runs RTA 2.11.3.0.

### Animal experiments

All animal procedures were approved by institutional ethic committee of CR Sainte-Justine Hospital. NOD/SCID/IL2Rγ null (NSG) mice were obtained from Humanized mouse platform of CR-CHU Saint-Justine. Mice were housed in the sterile animal facility under pathogen-free conditions. 5-week-old animals were separated for acclimatization and were injected with cells at 6 weeks of age. HeLa cells (shScrambled, shHCAR1b and KD+δS305A) were transduced with Renilla-Luciferase viral vectors and GFP-positive cells were sorted. Passage 3 of sorted cells were counted and 1 million cells were injected into animals. Mice were anesthetized with 2.5% isoflurane and cells were injected subcutaneously in the right flank after shaving and sterilizing the area for tumor growth monitoring, or injected into the tail vein for metastatic analysis of the cells. Two male and two female mice were used for each cell line. Animals were imaged once every 3 days for 5 weeks. Mice were anesthetized and injected with D-Luciferin (150mg/kg) 10 min before imaging. In vivo whole-body imaging was performed using Epi-Fluorescence and Trans-Fluorescence imaging system (OiS300, LabeoTech) and signal intensities were normalized and measured in radiance integrated density (photons ∣ s^−1^ ∣ sr^−1^ ∣ cm^−2^) using Fiji Macros. Animals were sacrificed after 5 weeks or at the study cut-off points (extreme abscess or 30% weight loss and morbid condition) and tumor and organs were harvested for histological analysis. *Immunohistochemistry:*

Tissue samples were fixed in 10% Formalin O/N at RT, and then embedded in paraffine. Paraffin embedded blocks were cut to 5μm sections and deparaffinized in Xylene and decreasing concentrations of ethanol. Antigen retrieval was done with Sodium Citrate buffer (10mM Sodium Citrate, 0.05% Tween, pH6) for 10 min with pressure cooker. Slides were washed with TBS / Triton X-100, and then blocked with 10% normal serum, 1% BSA in TBS for 2h at RT. Incubation with primary antibody was done ON in 4°C, and endogenous peroxidase activity was blocked with 0.3% H_2_O_2_. Secondary HRP-conjugated antibody was used for detection. Slides were developed with DAB reagent and counterstained with DAPI. Samples were dehydrated, mounted and visualized with Leica DMi8 wide-field microscope with monochrome color camera.

### Statistics

The number of samples per group, number of replicates and details of error bars are provided in the figure legends. Statistical tests were performed using GraphPad Prism 9.0 (GraphPad Software). For comparisons between two experimental groups, unpaired two-tailed t-tests were used, and for comparison of three and more groups Analysis of Variance (ANOVA) was used followed by Bonferroni post hoc correction test with * *P* < 0.05, ** *P* < 0.01, \*\*\**P* < 0.0001 significance levels. Data are shown as the mean□±□s.d, except data in panel fig. 7b, c & f which are mean ± s.e.m. Every dataset is composed of at least n≥3 independent experiments. List of genes from high-throughput experiments were compared with Venny(Oliveros, 2007).

