## Supplementary figures and images for "Nuclear Location Bias of HCAR1 Drives Cancer Malignancy through Numerous Routes"

### Supplemental figures

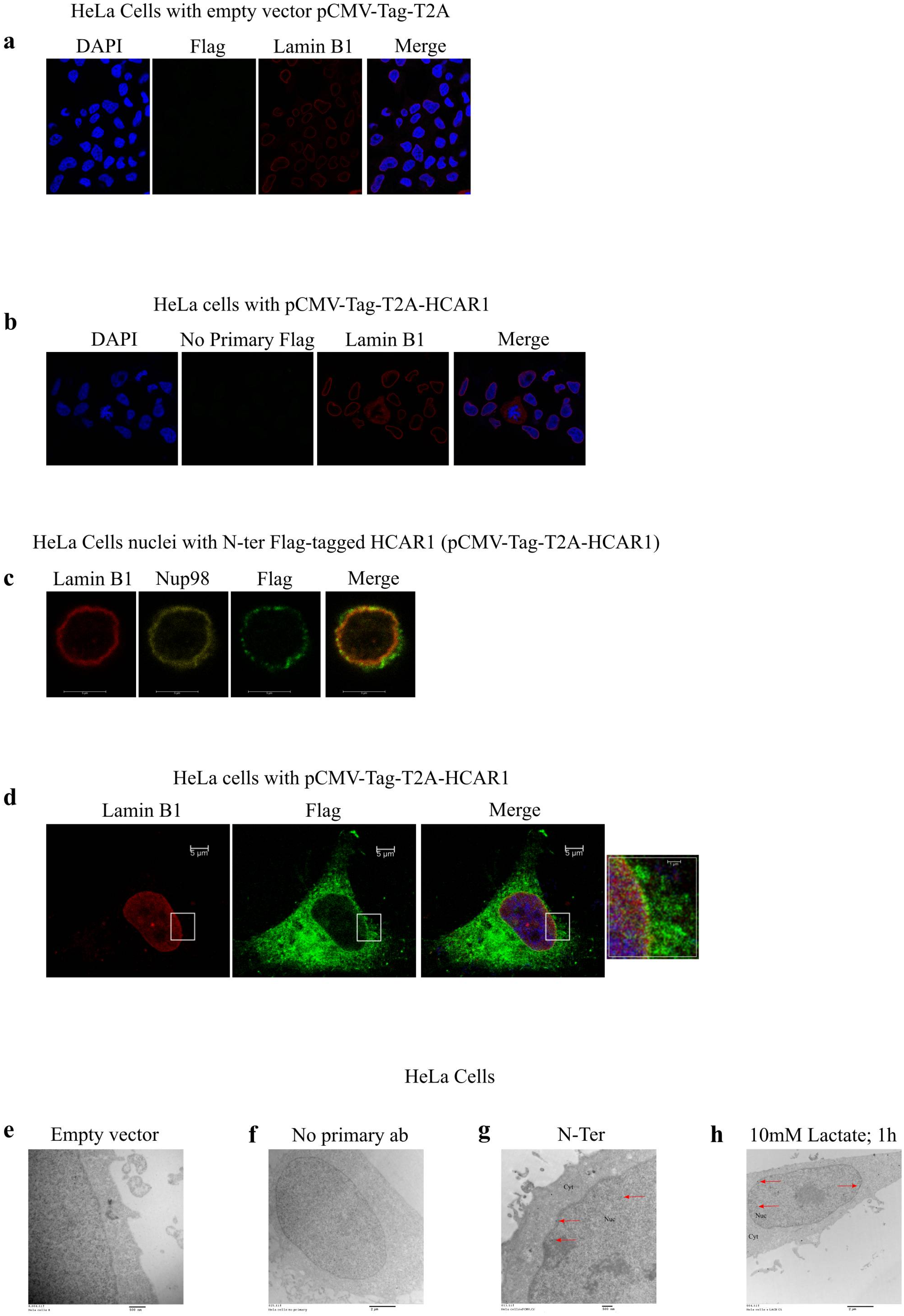

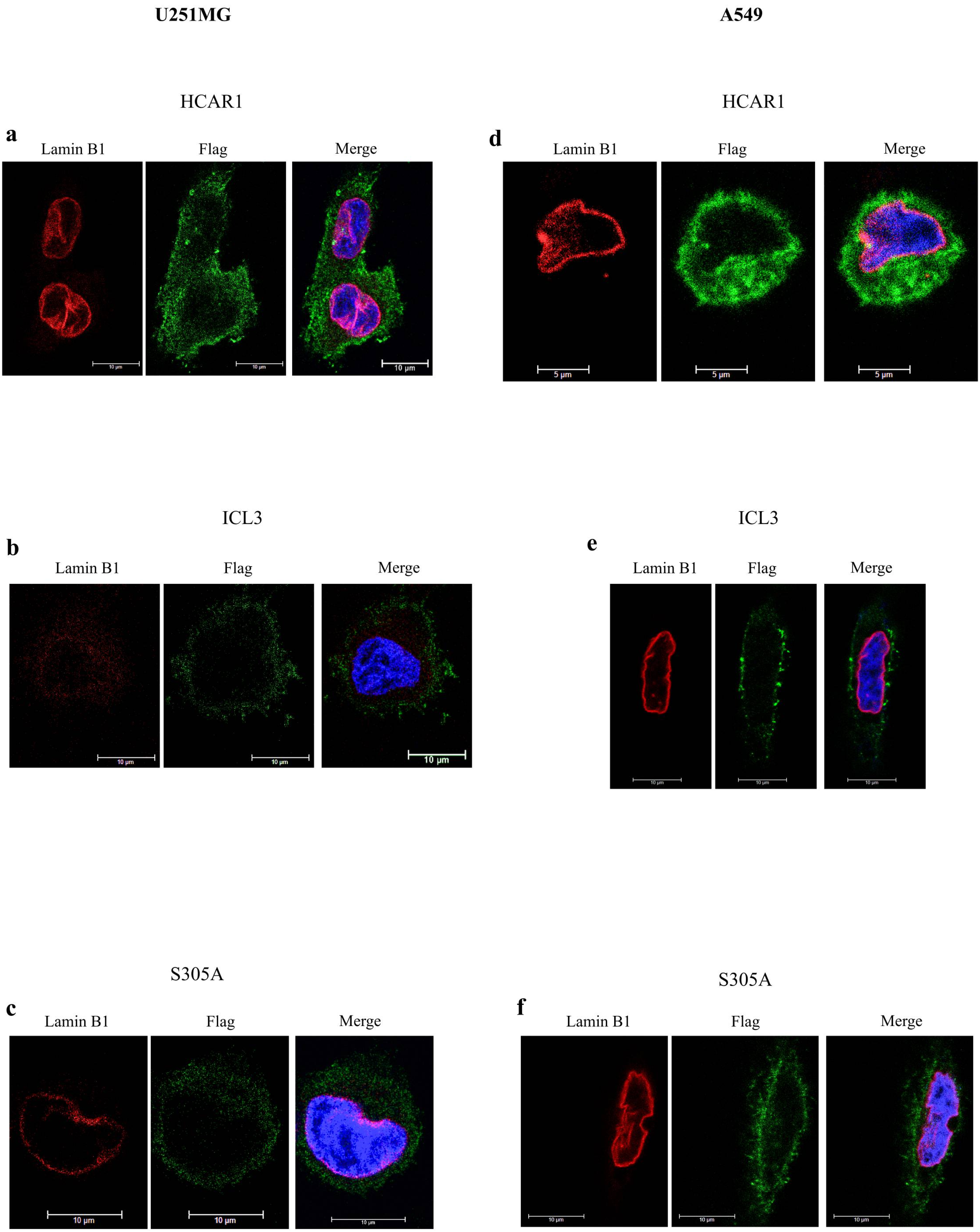

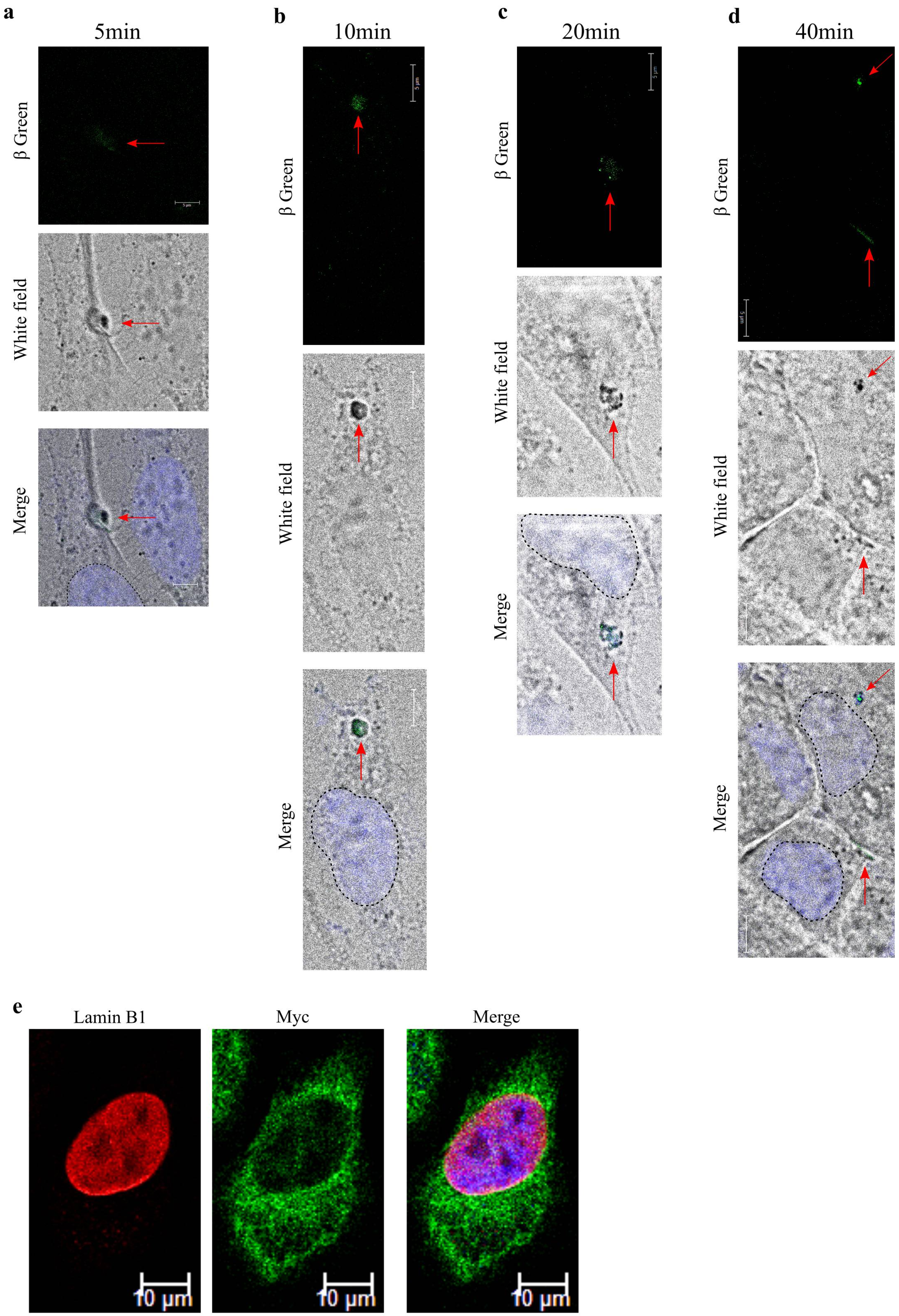

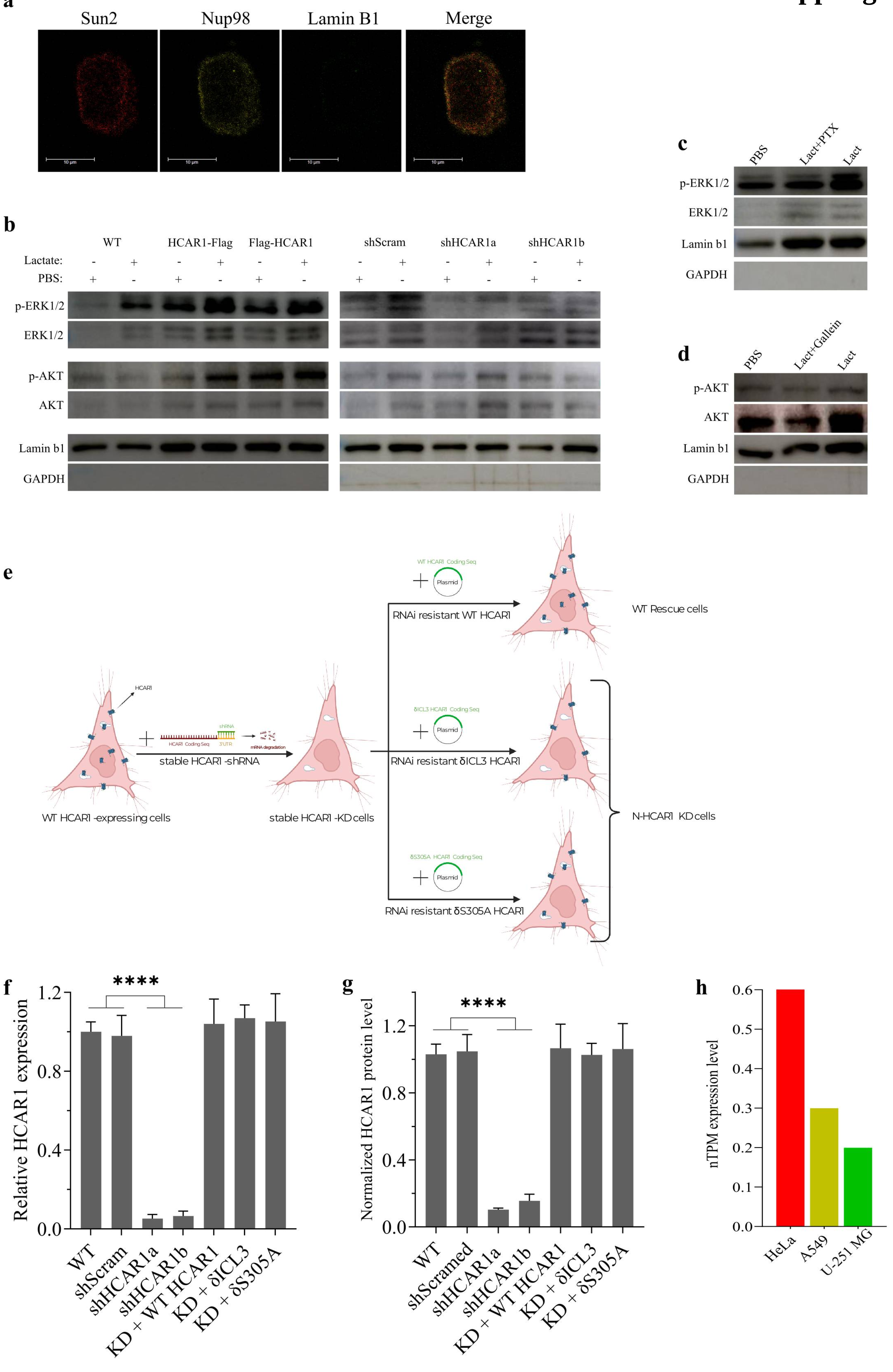

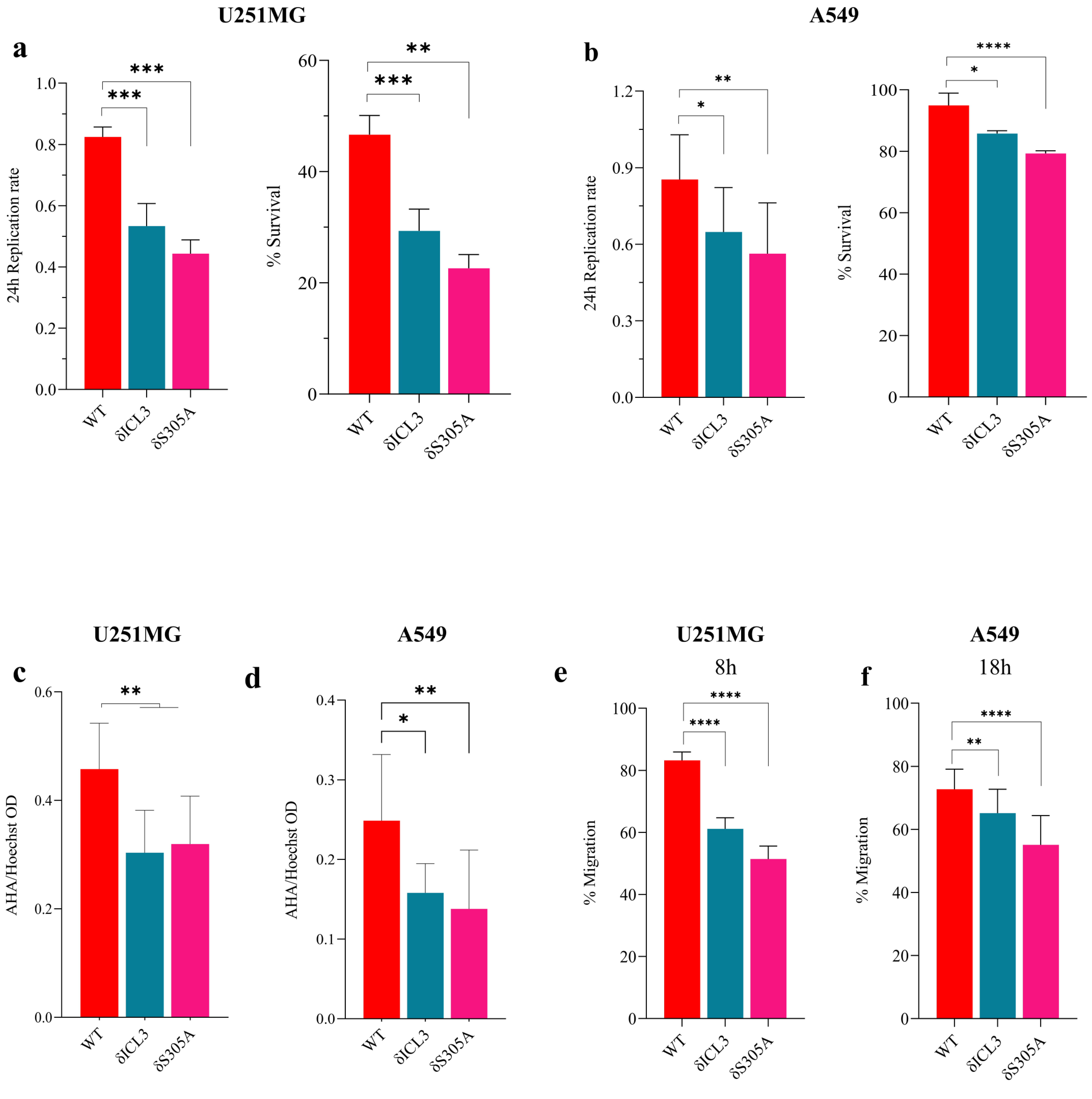

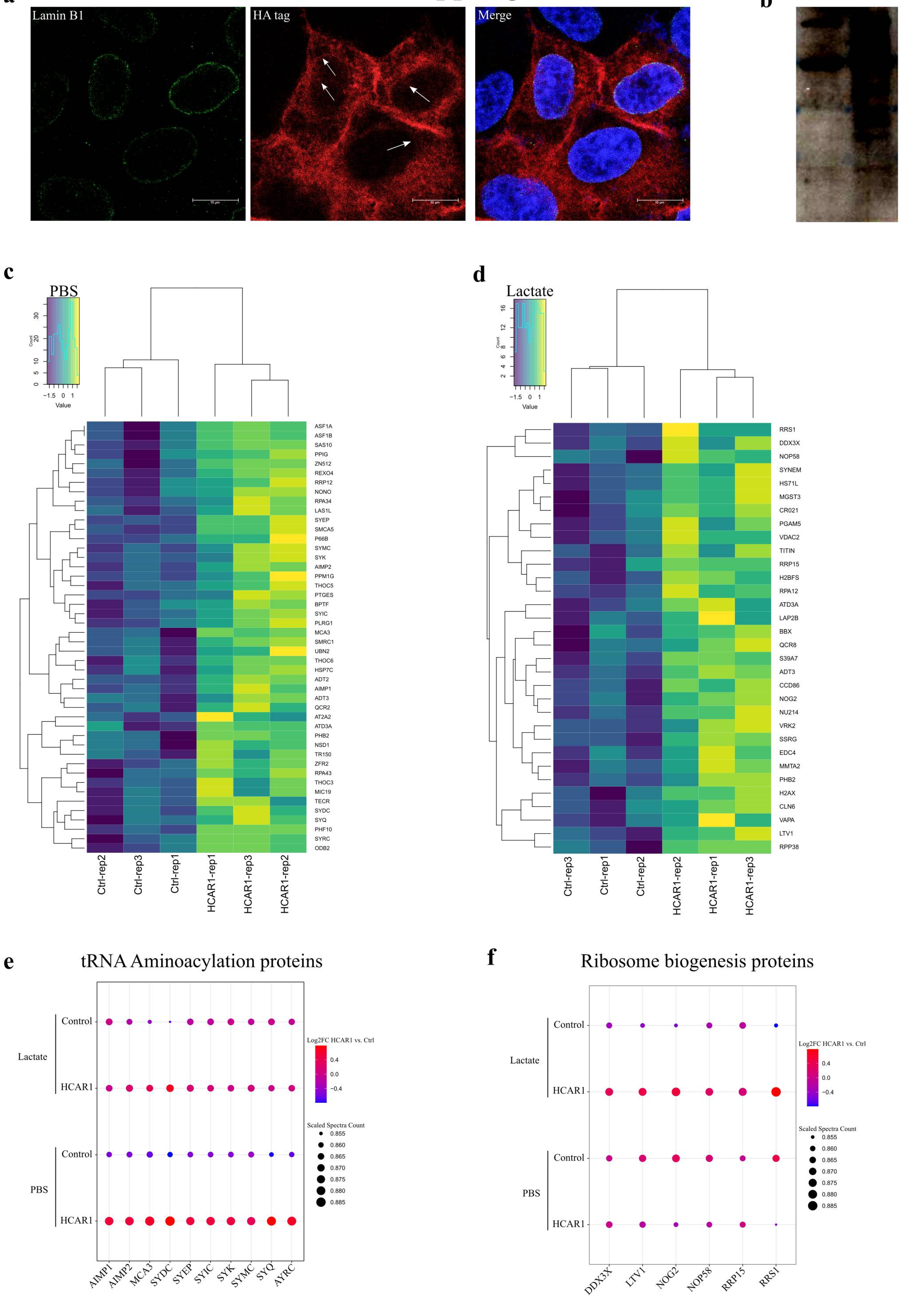

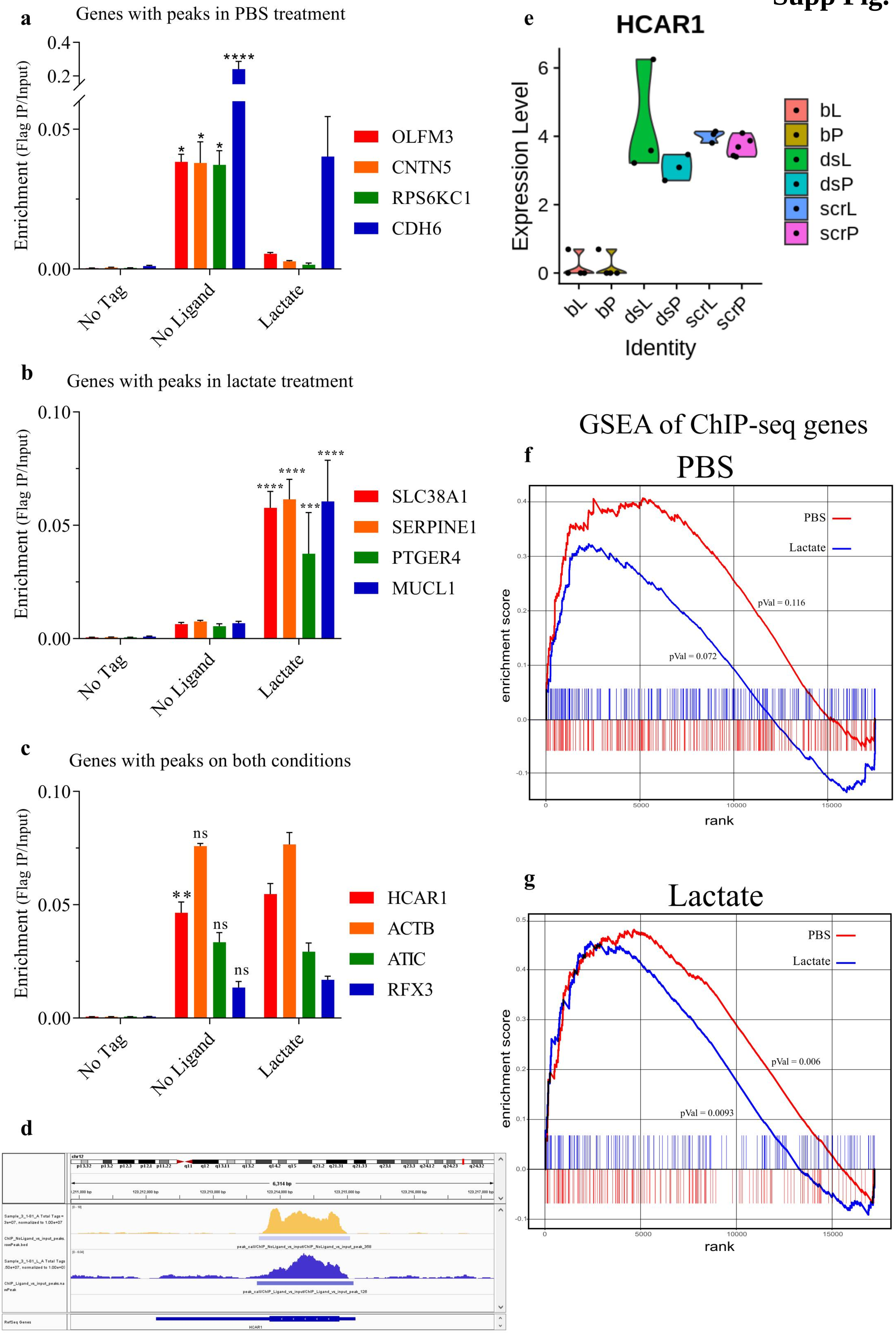
